# The heterogeneous landscape and early evolution of pathogen-associated CpG dinucleotides in SARS-CoV-2

**DOI:** 10.1101/2020.05.06.074039

**Authors:** Andrea Di Gioacchino, Petr Šulc, Anastassia V. Komarova, Benjamin D. Greenbaum, Rémi Monasson, Simona Cocco

**Affiliations:** Laboratoire de Physique de l’Ecole Normale Supérieure, PSL & CNRS UMR8063, Sorbonne Université, Université de Paris, F-75005 Paris, France; School of Molecular Sciences and Center for Molecular Design and Biomimetics, The Biodesign Institute, Arizona State University, 1001 South McAllister Avenue, Tempe, Arizona 85281, USA; Molecular Genetics of RNA viruses, Department of Virology, Institut Pasteur, CNRS UMR-3569, 75015 Paris, France; Computational Oncology, Department of Epidemiology and Biostatistics, Memorial Sloan Kettering Cancer Center, 1275 York Avenue New York, NY 10065

**Keywords:** ssRNA viruses, SARS-CoV-2, pathogen-associated molecular patterns (PAMPs), pattern recognition receptors (PRRs), viral host mimicry, CpG motifs, evolution of synonymous mutations

## Abstract

COVID-19 can lead to acute respiratory syndrome, which can be due to dysregulated immune signaling. We analyze the distribution of CpG dinucleotides, a pathogen-associated molecular pattern, in the SARS-CoV-2 genome. We find that the CpG content, which we characterize by a force parameter that accounts for statistical constraints acting on the genome at the nucleotidic and amino-acid levels, is, on average, low compared to other pathogenic betacoronaviruses. However, the CpG force widely fluctuates along the genome, with a particularly low value, comparable to the circulating seasonal HKU1, in the spike coding region and a greater value, comparable to SARS and MERS, in the highly expressed nucleocapside coding region (N ORF), whose transcripts are relatively abundant in the cytoplasm of infected cells and present in the 3’UTRs of all subgenomic RNA. This dual nature of CpG content could confer to SARS-CoV-2 the ability to avoid triggering pattern recognition receptors upon entry, while eliciting a stronger response during replication. We then investigate the evolution of synonymous mutations since the outbreak of the COVID-19 pandemic, finding a signature of CpG loss in regions with a greater CpG force. Sequence motifs preceding the CpG-loss-associated loci in the N ORF match recently identified binding patterns of the Zinc finger Anti-viral Protein. Using a model of the viral gene evolution under human host pressure, we find that synonymous mutations seem driven in the SARS-CoV-2 genome, and particularly in the N ORF, by the viral codon bias, the transition-transversion bias and the pressure to lower CpG content.

## 1 Introduction

When a virus enters a new host, it can present pathogen-associated molecular patterns (PAMPs) that are rarely seen in circulating strains that have adapted to that host’s immune environment over evolutionary timescales. The emergence of SARS-CoV-2, therefore, provides a rare window into innate immune signaling that may be relevant for understanding immune-mediated pathologies of SARS-CoV-2, anti-viral treatment strategies, and the evolutionary dynamics of the virus, where evidence for selective pressures on viral features can reflect what defines “self” in its new host. As a case in point, the 1918 influenza pandemic was likely caused by a strain that originated in water fowl and entered the human population after possible evolution in an intermediate host. That viral genome presented CpG dinucleotides within a context and level of density rarely found in the human genome where they are severely underrepresented, particularly in a set of genes coding for the proteins associated with antiviral innate immunity [1, 2, 3, 4]. Over the past century the 1918 H1N1 lineage evolved in a directed manner to lower these motifs and gain UpA motifs, in a way that could not be explained by its usage of amino-acid codon bias [4, 5]. It has since been found that these motifs can engage the pattern recognition receptors (PRRs) of the innate immune system [6, 7], and directly bind the Zinc finger Anti-viral Protein (ZAP) in a CpG-dependent manner [8, 9, 10, 11]. Hence, the interrogation of emergent viruses from this perspective can predict novel host virus interactions.

COVID-19 presents, thus far, a different pathology than that associated with the 1918 H1N1, which was disproportionately fatal in healthy young adults. It has been characterized by a large heterogeneity in the immune response to the virus [12, 13, 14] and likely dysregulated type-I interferon (IFN) signaling [15, 16, 17, 18]. Various treatments to attenuate inflammatory responses have been proposed and are currently under analysis or being clinically tested [19, 20]. It is therefore essential to quantify pathogen-associated patterns in the SARS-CoV-2 genome for multiple reasons. The first is to better understand the pathways engaged by innate immune system and the specific agonists to help build better antiviral therapies. Another is to better predict the evolution of motif content in synonymous mutations in SARS-CoV-2, as it will help understand the process and timescales of attenuation in humans. Third is to offer a principled approach for optimizing vaccine strategy for designed strains [21, 22] to better reflect human-genome features.

In this work we will use the computational framework developed in [4] to carry out a study of non-self associated dinucleotide usage in SARS-CoV-2 genomes. The statistical physics framework is based on the idea of identifying the abundance or scarcity of dinucleotides given their expected usage based on host features. It generalizes the standard dinucleotide relative abundance introduced in [1], as it can easily incorporate constraints in coding regions coming from amino-acid content and codon usage. The outcome of the approach are selective forces [4] that characterize the deviations with respect to the number of dinucleotides which is statistically expected under a set of various constraints. Such formalism has further been applied to identify non-coding RNA from repetitive elements in the human genome expressed in cancer that engage PRRs [23], to characterize the CpG evolution through synonymous mutations in H1N1 [4], and to characterize local and non-local forces on dinucleotides across RNA viruses [7].

We perform an analysis of the landscape of CpG motifs and associated selective forces in SARS-CoV-2 in comparison with other ssRNA viruses and other genomes in the coronavirus family in order to understand specific PAMP features in the new SARS-CoV-2 strains. We also focus on the heterogeneity of CpG motif usage along the SARS-CoV-2 genome (Sec. 2.2 and Sec. 2.3). Finally we use a model of viral genome evolution under human host pressure from the CpG force to study synonymous mutations, and in particular those which change CpG content, observed since the SARS-CoV-2 entered the human population (Sec. 2.4), and study sequence motifs preceding CpG loss loci (Sec. 2.5). The model is used to score all possible synonymous mutations from an ancestral sequence sampled in Wuhan at the beginning of the COVID-19 pandemic (GISAID ID: EPI ISL 406798) and is validated on single nucleotide variants observed in the sequence data collected so far (Sec. 2.6). This approach points out at hotspots where new mutations will likely attenuate the virus, while evolving in contact with the human host.

## 2 Results

### 2.1 Definition of coding and non-coding CpG forces

To characterize CpG dinucleotide usage on SARS-CoV-2 genome we have computed the CpG forces following the approach introduced in [4] and described in Methods (4.1, 4.2). CpG forces are intensive parameters that characterize the abundance or scarcity of CpG dinucleotides in a DNA or RNA sequence with respect to their expected usage relative to a reference nucleotide distribution. We propose two frameworks to define these reference distribution, schematically represented in Fig. 1. In the non-coding framework, nucleotides are randomly drawn according to a fixed nucleotide bias, while in the coding framework the amino-acid sequence is fixed, and the codon bias defines the reference distribution.

**Figure 1:**
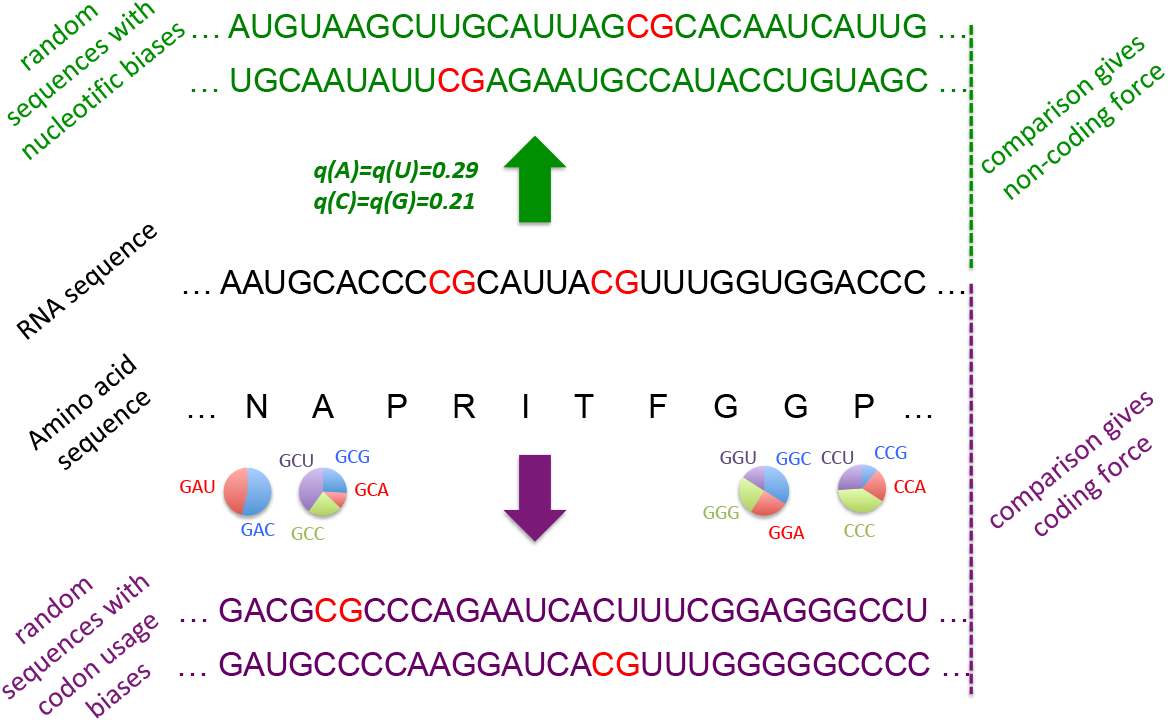
Schematic definitions of CpG non-coding and coding forces. The natural DNA (or RNA) sequence to be analyzed is shown in black, with CpG dinucleotide motifs in red. The force is computed by comparing the occurrences number of CpG with ensembles of random sequences fulfilling some of the constraints acting on the natural sequence, see Methods for details. Top, *non-coding* framework, in green: sequences of the same lengths can be generated by randomly drawing nucleotides according to prescribed frequencies, here taken from the human genome. Bottom: when the sequence under consideration codes for a protein (sequence of amino acids in black letters), random sequences (violet letters) can be generated in a *coding* framework as follows. For each amino acid, a licit codon is randomly chosen according to prescribed codon usage (here, computed from the coding regions in the human genome). Notice that the above computational frameworks are not restricted to CpG, and can be applied to other dinucleotidic motifs.

#### Non-coding forces

The CpG non-coding force relative to the sequence nucleotide bias essentially captures the same information as the relative abundance index, 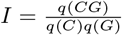, where *q*(*CG*), *q*(*C*), *q*(*G*) are, respectively, the frequencies of CpG dinucleotides and of *C*, *G* nucleotides in the sequence [1]. The CpG non-coding force is well approximated by the logarithm of the relative abundance index: *f ≈* log *I* (see Suppl. Sec. SI.3 and Fig. SI.12). Positive and negative forces correspond therefore to, respectively, excess and scarcity of dinucleotides with respect to their expected occurrences determined by the nucleotide bias.

Table 1 shows that the CpG non-coding forces for human coding [4] and non-coding RNA [23] (relative to the human nucleotide bias) are negative, and particularly low for type-I IFN transcripts involved in the innate immune system [4], confirming that CpG motifs are overall scarce in the human genome [1, 2, 3, 4]. As shown in Table 1 strongly pathogenic viruses in humans, such as Ebola, the Spanish flu H1N1 (1918) and the avian flu H5N1 (2005), are characterized by large CpG forces compared to the average force on human RNAs. The CpG forces value can then be used as an indicator for the propensity of a viral sequence to be sensed by PRRs as non-self and engage the human innate immune response [4, 7, 23]. A comparative analysis for non-coding force in the *Coronaviridae* family will be discussed in the following Sections.

**Table 1:**
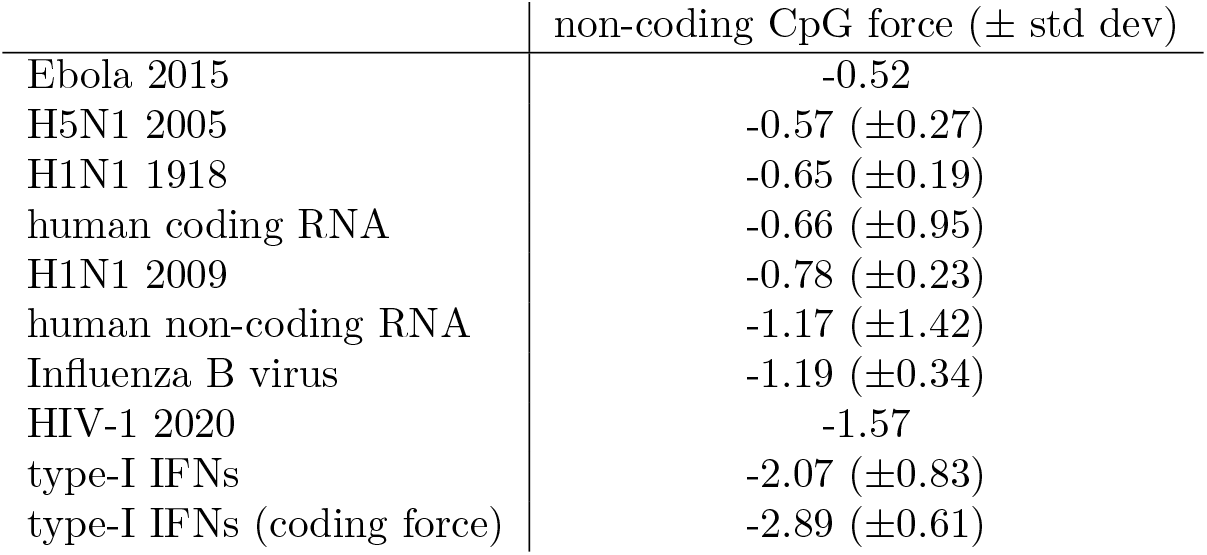
Global non-coding CpG forces for some ssRNA viruses, compared to human RNAs. The distribution of forces are computed from all genomic segments and their averages and standard deviations are given (with segment contribution weighted by segment length). All forces are computed with respect to human nucleotide bias. Data used: human cDNA and ncRNA as annotated in HG38 assembly, transcripts coding for type-I IFN’s genes as annotated as annotated in HG38. Viral ssRNAs were obtained from NCBI [24] Virus database (strains used: H5N1: A/Anhui/1/2005, H1N1: A/Aalborg/INS132/2009 and A/Brevig Mission/1/1918, Ebola: COD/1995/Kikwit-9510623, Influenza B virus: B/Massachusetts/07/2020, HIV-1: HK JIDLNBL S003).

#### Coding forces

The CpG coding force is based on the comparison of CpG occurrences in a coding RNA (or DNA) sequence and random synonymous sequences (associated to the same amino acids) drawn according to prescribed codon usage, cf. Figure 1. The computation of coding forces relative to the human codon usage for SARS-CoV-2 will be discussed in Sec. 2.3 and in Sec. 2.4 it will be used to characterize the evolution of SARS-CoV-2 sequences through synonymous mutations, under the pressure of the human host. To allow for easy comparison and later access we list in Table 2 all CpG coding forces for the *Coronaviridae* family, as well as their non-coding counterparts.

**Table 2:**
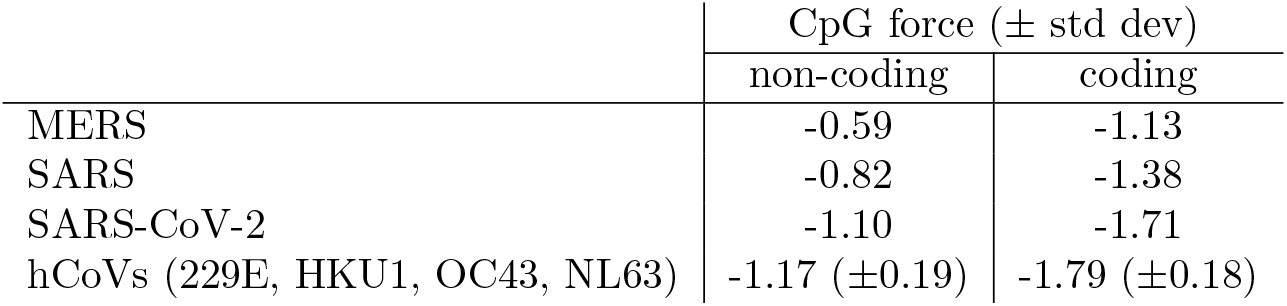
Global non-coding and coding CpG forces for *Coronaviridae* family viruses. All forces are computed with respect to human nucleotide (non-coding forces) or codon bias (coding forces).

### 2.2 The landscape of CpG forces in SARS-CoV-2 is strongly heterogeneous

We first computed the global non-coding force on CpG dinucleotides for SARS-CoV-2, a variety of other ssRNA viruses, and other viruses from *Coronaviridae* family affecting humans or other mammals (Fig. 2a).

**Figure 2:**
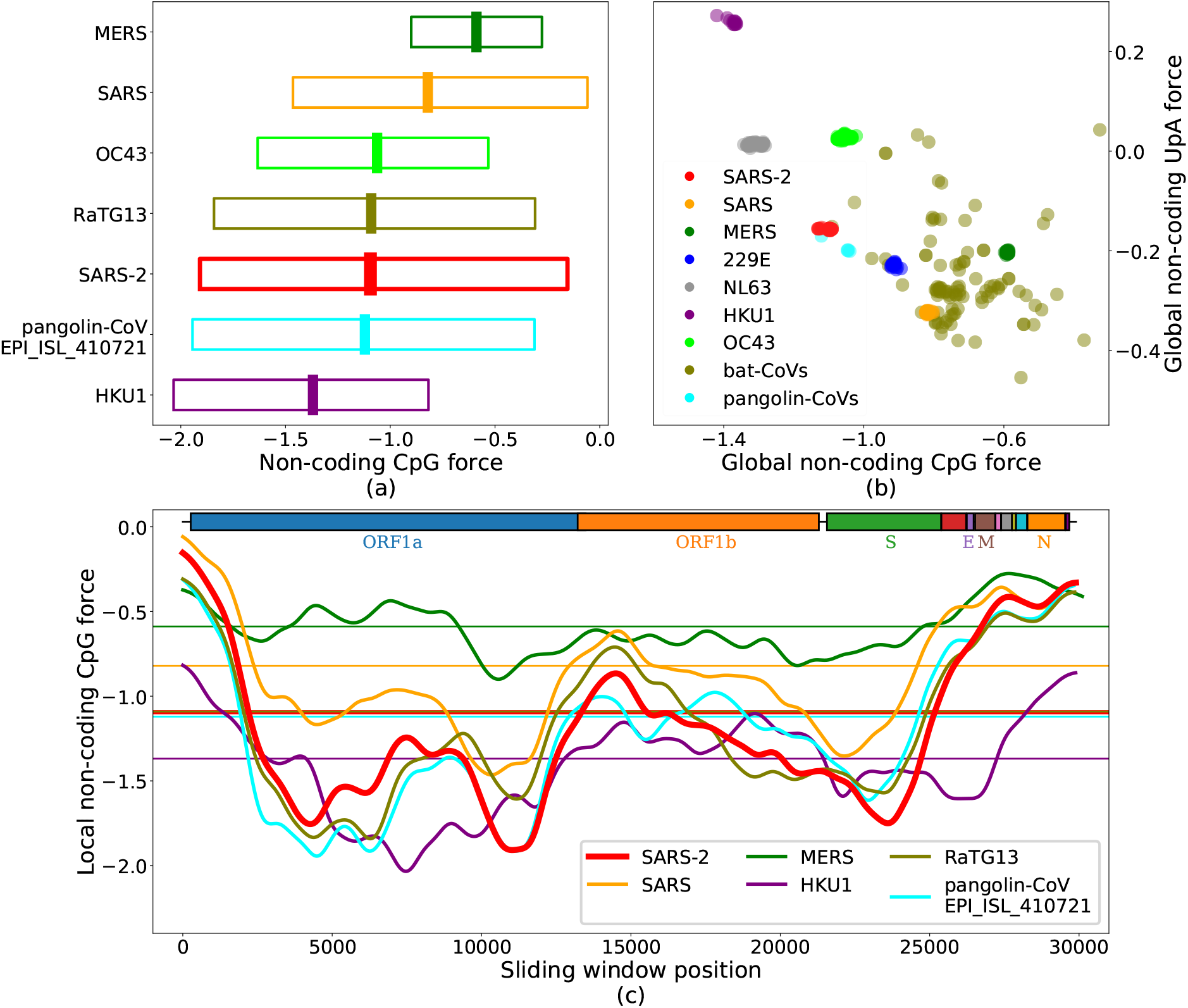
CpG and UpA non-coding forces, and local fluctuations in the genomes of the *Coronaviridae* family. (a): Non-coding forces on CpG dinucleotides for SARS-CoV-2 and other coronaviruses. The central thick lines show global forces over the whole genomes, while bars span from the minimal to the maximal values of the local forces computed on sliding windows along the sequence of 3kb (narrowed up to 1.5 kb at the edges), smoothed through a Gaussian average. (b): UpA vs. CpG global forces. Species are well clustered, due to the large similarity of the sampled sequences, except for viruses circulating in bats composed of several diverse strains. Notice the anticorrelation between CpG and UpA forces (Pearson *r* = −0.69, with a p-value of ≃ 2.10^−77^, and *r*^2^ = 0.48). (c): Local CpG force analysis in sliding windows of 3 kb along the genome for some coronaviruses, smoothed through a Gaussian average; Horizontal lines correspond to the global forces shown in panel (b). Boxes on top of the panel show protein-coding domains. The force is highly variable along the genome, with much larger values in certain regions (such as the N ORF) than in others (e. g. S ORF). The maximum value of the local CpG force hints at the similarity of SARS-CoV-2 with the most pathogenic viruses, see Table 2. Data from VIPR [31] and GISAID [32], see Methods Sec. 4.6 and Suppl. Sec. SI.1 for details on data analysis.

The value ≃ −1.1 of the global non-coding force for SARS-CoV-2 is comparable to the one for human non-coding RNA and lower than other strongly pathogenic viruses in humans, such as H1N1, H5N1, Ebola (see Table 1). Among the coronaviruses (see Fig. 2a and Table 2) MERS shows the highest CpG force followed by SARS, while some bat coronaviruses have even strongest CpG force. SARS-CoV-2 is among the viruses with lower global CpG force; its value is median among the hCoV that circulate in humans with less pathogenicity, between HKU1 with a smaller CpG force and OC43 with a larger one (See Suppl. Fig. SI.4a for a comparison with other hCOVs). The absence of a straightforward correlation between global CpG force and the pathology of a coronavirus in humans calls for a finer, local analysis of CpG forces we report below.

Figure 2b compares the forces^1^ acting on CpG and UpA motifs within the *Coronaviridae* family, with a particular emphasis on the genera *Alphacoronavirus* and *Betacoronavirus*, and on those viruses which infect humans [25]; for other dinucleotides, see Suppl. Fig. SI.1. We observe an anti-correlation between UpA and CpG forces (Correlation Coefficient R-Squared *r*^2^ = 0.48). UpA is the CpG complementary motif corresponding to the two nucleotidic substitutions more likely to occur in terms of mutations, as transitions have larger probability with respect to transversions and are less likely to result in amino-acid substitutions. Such anti-correlations are not observed with motifs that are one mutation away from CpG (*r*^2^ = 0.2 for CpA versus CpG and *r*^2^ = 0.01 for UpG versus CpG, Suppl. Fig. SI.3).

To go beyond the global analysis we study the local non-coding forces acting on CpG in fixed-length windows along the genome. Results for SARS, MERS, SARS-CoV-2, hCoV-HKU1 and two representative sequences of bat and pangolin coronaviruses, chosen for their closeness to SARS-CoV-2, are reported in Fig. 2c. SARS-CoV-2 shows a strong heterogeneity of CpG forces along its genome: in some genomic regions, especially at the 5’ and 3’ extremities, SARS-CoV-2, SARS and MERS (together with the bat and pangolin viruses) have a peak in CpG forces, which is absent in the hCoV-HKU1 (as well as in the other hCoVs, see Suppl. Fig. SI.4). The heterogeneous CpG content in SARS-CoV-2 has been also pointed out in [26].

The high CpG forces at the extremities could have an important effect on the activation of the immune response via sensing, as the life cycle of the virus is such that the initial and final part of the genome are those involved in the subgenomic transcription needed for viral replication [27, 28]. During the infection many more RNA fragments from these regions are present in the cytoplasm than from the other parts of the viral genome. Consequently, despite the relatively low CpG content of SARS-CoV-2 compared to other coronaviruses, there can be high concentrations of CpG-rich RNA due to the higher transcription of these regions.

The similarity between the high values of the maximum local forces of SARS-CoV-2 and those of SARS, bat and pangolin coronaviruses shown in Fig. 2a confirms this pattern: MERS and SARS, viruses that are likely less well adapted to a human host, have the highest local peaks in CpG content, followed by SARS-CoV-2 and then by seasonal strains that circulate in humans. It is interesting to notice that high and very high levels of proinflammatory cytokines/chemokines (such as IL-6 and TNF-*α*) have been observed in, respectively, SARS and MERS and, at times, SARS-CoV-2 infection [12, 18, 29, 30]. These results are qualitatively corroborated by the simpler analysis of CpG motif density (Suppl. Fig. SI.2).

### 2.3 Forces acting on coding regions widely vary across structural proteins

We now restrict our analysis to the coding regions of SARS-CoV-2 and, in particular, on two structural proteins, N (nucleocapside) and S (spike) [21, 33, 34]. As stressed in Sec. 2.1 the computation of the force in such coding regions account for the extra constraints on the amino-acid content and takes the human codon bias as reference background, see Methods Sec. 4.1 and Sec. 4.2.

The landscape of coding CpG forces with respect to the human codon bias is shown, restricted to the coding regions of SARS-CoV-2 genome, in Fig. 3a, and compared to the non-coding forces from Fig. 2, with respect to the human nucleotide bias (dashed red lines). The two curves are similar apart from a global shift towards lower forces for the coding forces. Notice that this shift essentially vanishes if the non-coding force is computed with respect to the nucleotide bias computed on human coding RNAs only [35] (Fig. 3a). Apart from this global shift, the strongly heterogeneous landscape of the CpG coding forces along the SARS-CoV-2 genome does not substantially differ from the findings of Fig. 2c. In particular the peak of high CpG density and force is still present at the 5’ and the 3’ ends of the genome, including the N ORF, the envelope E ORF and membrane glycoprotein M ORF regions. In the S ORF region the coding CpG force remains small. Detailed results for the S and N ORFs are shown in, respectively, Figs. 3b and 3c. These structural proteins are present and quite similar across the *Coronaviridae* family, and allow us to compare several strains of coronaviruses. In the S ORF, SARS-CoV-2 shows the lowest global CpG force among the human-infecting betacoronaviruses, see Fig. 3c. The CpG force is much higher for protein N in SARS-CoV-2, immediately below the level of SARS and above that of MERS, see comparison with human-infecting members of the *Coronaviridae* family presented in Fig. 3b. The comparative analysis of forces in the E ORF (Suppl. Fig. SI.5b) gives results similar to the N ORF, while smaller differences in CpG force among coronaviruses that circulate in humans are observed for the M ORF (Suppl. Fig. SI.5c).

**Figure 3:**
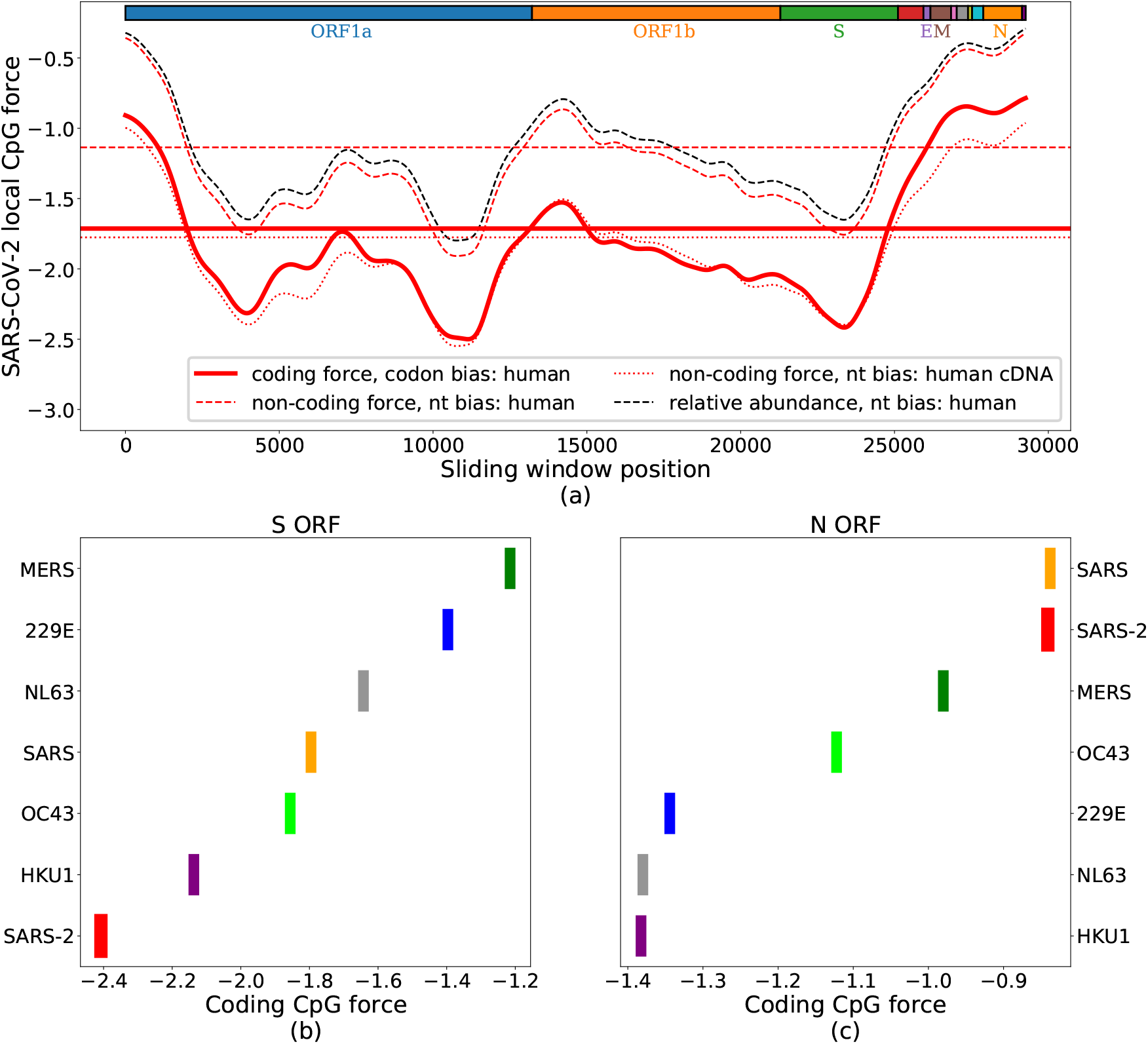
CpG local coding forces on SARS-CoV-2 coding regions. (a): Local forces, over sliding windows ranging of 3kb (narrowed up to 1.5kb at the edges), in the the coding regions of SARS-CoV-2 (pre-processed to ensure the correct reading frame). Full, thick red curve shows coding forces. The dashed and dotted lines show non-coding forces obtained by using nucleotide frequencies computed from, respectively, all human genome (including non coding RNA and coding RNA) and only coding human RNAs. Horizontal lines locate global forces. The black dashed line shows the relative abundance of CpG computed on the same sliding windows, with the same nucleotide frequency used for the dashed red line. Boxes on top of the panel show protein-coding domains. (b) and (c): global forces for structural proteins (S and N) in the *Coronaviridae* family. This values were averaged on 4 to 20 sequences from VIPR [31] and GISAID [32], see Methods Sec. 4.6 and Suppl. Sec. SI.1 for details on data analysis.

### 2.4 Force-based model accounts for early evolution of synonymous CpG-related mutations in SARS-CoV-2

We now assess the ability of our CpG force model to predict biases in the synonymous mutations already detectable across the few months of evolution following the first sequencing of SARS-CoV-2 (data from GISAID [32]; Wuhan ancestral strain has GISAID ID: EPI ISL 406798, collected in Wuhan on 26 Dec 2019; last updated sequence 29 Sept 2020; see Methods Sec. 4.6). Barring confounding effects, we expect that high-force regions, such as N ORF, will be driven by the host immune system pressure towards a lower number of CpG motifs. Other regions, such as S ORF, already have low CpG content and would feel no pressure to decrease the CpG content, so random mutation would likely leave their CpG number unaffected or increase it. We define in the following Synonymous Single Nucleotide Variants (Syn-SNV) as nucleotide synonymous substitution with respect to the Wuhan ancestral strain, observed at least in 5 collected sequences (0.01% of the sample)^2^.

Figure 4a (bottom and middle panels) shows that many Syn-SNV that decrease the number of CpG are located at the 5’ and 3’-end of the sequence, in correspondence with the high peak in CpG local force, notably in the N ORF region and at the 5’ extremity of the genome. Conversely Syn-SNV that increase the number of CpG are more uniformly spread along the sequence. The behaviors of the local CpG force and of the local density of CpG decreasing Syn-SNV, computed on the same sliding windows along the sequence, show strong similarities, see middle panels in Fig. 4a.

**Figure 4:**
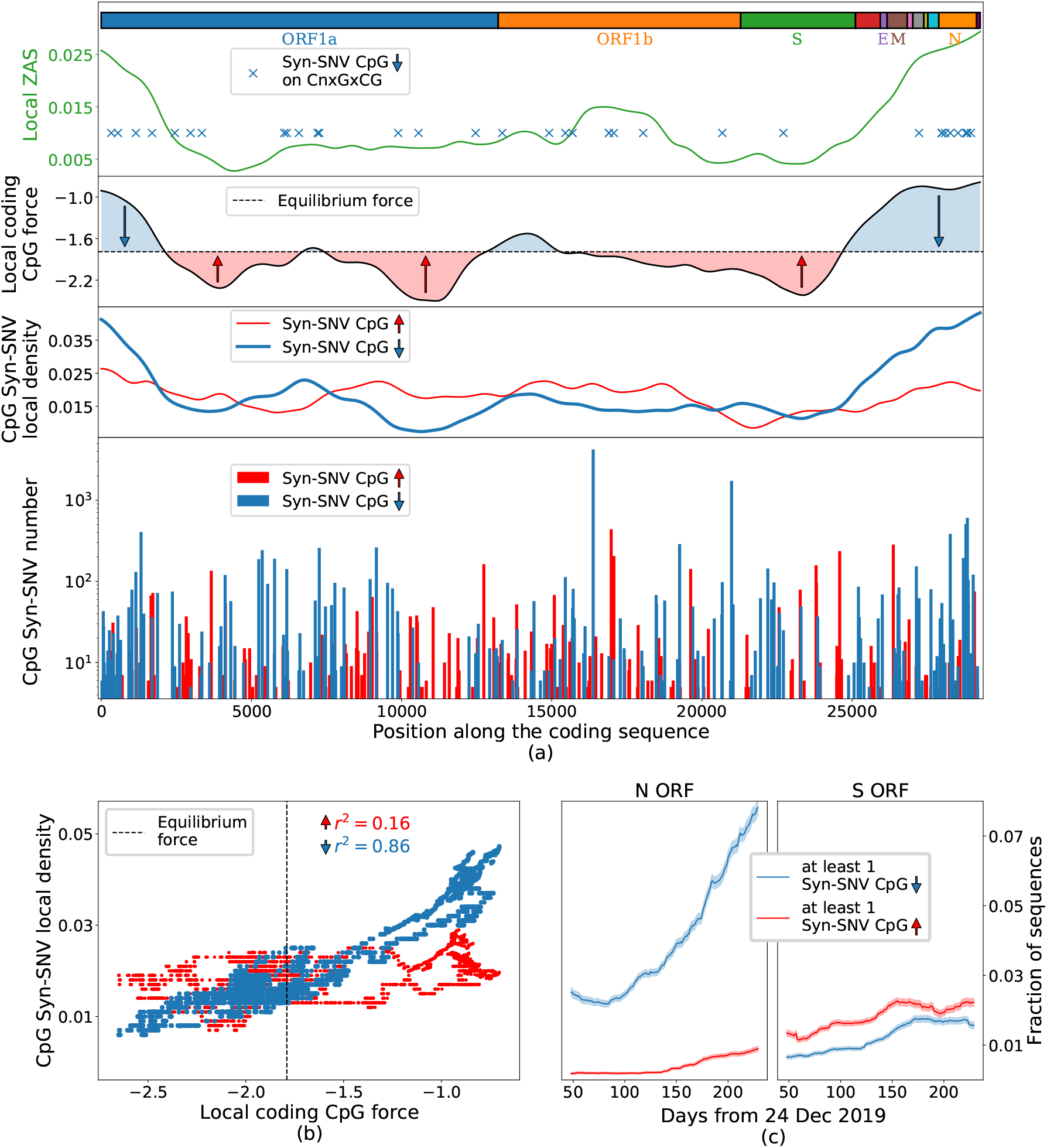
Analysis of synonymous mutations in the early evolution (up to October 2020) of SARS-CoV-2 genome, CpG drive and ZAP binding motifs. (a): Bottom: Counts of syn-SNV that increased (red) and decreased (blue) the CpG content. Middle: sliding average of syn-SNV increasing (red) and decreasing (blue) CpG with windows of 3 kb and a Gaussian smoothing; black line: local coding CpG force with same sliding average and smoothing, dashed black line: putative equilibrium force (−1.79) for SARS-CoV-2 coding regions. The area between the local CpG force and the equilibrium CpG force is filled in blue/red for local CpG force larger/smaller than the equilibrium one. Upper sub-panel: The local ZAP affinity score (ZAS), computed on sliding windows of 3 kb; blue crosses mark SNV removing CpG motifs in a CnxGxCG patterns. Boxes on top of the panel show protein-coding domains. (b): scatter plot of the local CpG force (black curve in panel (a), without smoothing procedure) versus the local density of CpG decreasing (blue points) or increasing (red points) Syn-SNV. Dashed vertical line: putative equilibrium force. (c): fraction of sequences in the data with at least one Syn-SNV decreasing (blue curve) or increasing (red curve) the CpG content in the N ORF (left) and S ORF (right), as function of time. To reduce noise, for each point we considered all the sequences collected in a temporal sliding window of 100 days centered on the point. Data from GISAID [32], see Methods Sec. 4.6 (last update 05 Oct 2020) for details on data analysis. Ancestral genome: GISAID ID: EPI ISL 406798 (Wuhan, 26-12-2019).

To better explain this CpG mutational trend along the sequence we define a putative equilibrium CpG force of the SARS-CoV-2 genome in human host, as the average CpG force of hCoVs in Table 2: hCoVs have long been circulated in humans and are, therefore, supposed to be close to equilibrium with their host. Other choices for equilibrium force will be discussed later. Regions with a CpG force much larger than the equilibrium one are predicted to be under strong evolutionary pressure to decrease their CpG content. This prediction is confirmed by the fact that CpG– decreasing syn-SNV are much more frequent than CpG–increasing ones, see Fig. 4a, middle. Conversely, in regions with forces slightly smaller than the equilibrium force value, the presence of a small evolutionary pressure to increase CpG is confirmed by the fact that CpG–increasing syn-SNV are slightly more frequent than CpG–decreasing ones. The scatter plot of the local forces and densities of CpG–increasing (blue) and –decreasing (red) syn-SNV along the SARS-CoV-2 Wuhan ancestral strain is shown in Fig. 4b. The correlation coefficient is much larger for CpG– decreasing syn-SNV (*r*^2^ = 0.85) than for CpG–increasing mutations (*r*^2^ = 0.16). The two scatter plots cross at a local CpG force *f* ≃ 1.8 ± 0.2, very close to the equilibrium force, *f_eq_* = −1.79. This result supports our choice of the equilibrium force. The global force of SARS-Cov-2 (*f* = −1.71) is also compatible with this crossing point. On the contrary, other possible choices for the equilibrium force, such as the coding force computed on human type-I IFNs, *f* = −2.89, would not match the crossing point. The results above suggest to introduce, as will be done in Sec. 2.6, the CpG drive defined as the difference between the CpG local force and the equilibrium CpG force. Table 3 complements Fig. 4 with a detailed description of CpG–decreasing and –increasing Syn-SNV along the ORFs and the 5’ and 3’ untranslated regions (UTRs) of SARS-CoV-2 genome. The regions with high negative CpG drive have a large density of CpG removing mutations, see for instance 5’UTR, 3’UTR, N ORF, and M ORF. Importantly, Syn-SNV are in many loci across the sequence (Fig. 4a), and taking into account Syn-SNV counts in the sequence data or considering unique Syn-SNV does not qualitatively affect our conclusions. Focusing on N ORF, remarkably 21% (47% with count) of Syn-SNV remove a CpG motif. Such percentage represents a fraction of 75% (94% with counts), among the total number of Syn-SNV affecting a CpG. On the opposite, the regions with a small negative or positive drive such as ORF1a, ORF1b and S ORF have a low density of CpG affecting mutations and among Syn-SNV affecting CpG motifs the percentage for Syn-SNV adding a CpG motif or removing a CpG motif are comparable. For S ORF, having the largest positive drive, the large majority of synonymous variants, 85% (92% with counts), leaves the CpG content unchanged with only few, 7% (4% with counts), Syn-SNV affecting a CpG motif. Among Syn-SNV affecting CpG, a slight predominance of CpG increasing Syn-SNV is observed with 53% (56% with counts) CpG increasing against 47% (44% with counts) CpG decreasing Syn-SNV. Last of all, Fig. 4c shows that, in N ORF, a rapid accumulation of CpG removing Syn-SNV is observed in the sampled sequences as a function of the delay between the time of collection and the beginning of the COVID-19 pandemic. This increase is much steeper that the gradual rise of Syn-SNV increasing CpG occurrences. In the S region, on the contrary, a gradual rise of Syn-SNV is observed both for CpG–increasing and –decreasing mutations, with a slight predominance of CpG–increasing ones.

**Table 3:** CpG drives and analysis of synonymous SNV changing CpG along the SARS-CoV-2 genome. The table gives, for all the ORFs and the 5’ and 3’ UTRs of SARS-CoV-2 ancestral genome, the length of the region (L), the number of CpG motifs (CpG), the CpG drive (*f_eq_* − *f*), the syn-SNV and the total numbers and percentages of syn-SNV removing a CpG motif (CG) or adding it (CG), with respect to total number of syn-SNV (/SNV) or to the total number of syn-SNV affecting CpG (/CpGSNV). For the non-coding 5’ and 3’-UTRs all SNV are taken into account with no restriction to syn-SNV and the non-coding forces are used; the equilibrium force is −1.16 (and not −1.79 as for ORFs) releasing such constraint. UTRs and ORFs and are sorted according to the density of CpG removing SNV (CG SNV/L). The regions underlined in bold are the most reliable for statistical analysis as they present at least 20 syn-SNV. Numbers and percentages of SNV are given with and without taking into account SNV counts. Data from GISAID [32], see Methods Sec. 4.6 for details on data analysis (last update 05 Oct 2020). Ancestral genome GISAID ID: EPI ISL 406798. SNV with less than 5 counts are excluded from the data.

|  | L | CpG | CpG drive | SNV | CpG↓SNV |  |  | CpG↑SNV |  |  |
| --- | --- | --- | --- | --- | --- | --- | --- | --- | --- | --- |
|  |  |  |  |  | tot | /SNV | /CpG↓SNV | tot | /SNV | /CpG↑SNV |
| **3'-UTR** | 162 | 5 | -0.87 | 81 | 18 | 22% | 90% | 2 | 2% | 10% |
|  |  |  | with counts: | 6020 | 1597 | 27% | 99% | 20 | 0.3% | 1% |
| **5'-UTR** | 211 | 13 | -1.67 | 56 | 19 | 34% | 95% | 1 | 2% | 5% |
|  |  |  | with counts: | 48806 | 47446 | 97% | 99% | 328 | 1% | 1% |
| **N** | 1260 | 39 | -0.95 | 115 | 24 | 21% | 75% | 8 | 7% | 25% |
|  |  |  | with counts: | 4745 | 2239 | 47% | 94% | 146 | 3% | 6% |
| **M** | 669 | 20 | -1.00 | 58 | 12 | 21% | 86% | 2 | 4% | 14% |
|  |  |  | with counts: | 5508 | 245 | 5% | 45% | 302 | 6% | 55% |
| ORF10 | 117 | 5 | -2.01 | 6 | 2 | 33% | 100% | 0 | 0% | 0% |
|  |  |  | with counts: | 233 | 14 | 60% | 100% | 0 | 0% | 0% |
| **ORF7a** | 366 | 7 | -0.61 | 23 | 5 | 22% | 56% | 4 | 17% | 44% |
|  |  |  | with counts: | 544 | 243 | 45% | 88% | 33 | 6% | 12% |
| **ORF8** | 366 | 8 | -0.82 | 25 | 5 | 20% | 63% | 3 | 12% | 38% |
|  |  |  | with counts: | 2146 | 107 | 5% | 86% | 17 | 1% | 14% |
| **ORF3a** | 828 | 17 | -0.62 | 54 | 10 | 19% | 67% | 5 | 9% | 33% |
|  |  |  | with counts: | 1530 | 244 | 16% | 87% | 36 | 2% | 13% |
| **ORF1a** | 13203 | 160 | 0.05 | 848 | 86 | 10% | 52% | 81 | 10% | 49% |
|  |  |  | with counts: | 79937 | 3930 | 5% | 72% | 1560 | 2% | 28% |
| **ORF1b** | 8088 | 115 | 0.003 | 432 | 47 | 11% | 48% | 52 | 12% | 53% |
|  |  |  | with counts: | 33788 | 7342 | 22% | 84% | 1434 | 4% | 26% |
| E | 228 | 11 | -1.84 | 10 | 1 | 10% | 100% | 0 | 0% | 0% |
|  |  |  | with counts: | 311 | 69 | 22% | 100% | 0 | 0% | 0% |
| **S** | 3822 | 29 | 0.61 | 223 | 16 | 7% | 47% | 18 | 8% | 53% |
|  |  |  | with counts: | 14540 | 571 | 4% | 44% | 729 | 5% | 56% |
| ORF6 | 186 | 1 | 0.18 | 10 | 0 | 0% | 0% | 2 | 20% | 100% |
|  |  |  | with counts: | 275 | 0 | 0% | 0% | 40 | 15% | 100% |
| ORF7b | 132 | 1 | -0.28 | 9 | 0 | 0% | 0% | 2 | 22% | 100% |
|  |  |  | with counts: | 406 | 0 | 0% | 0% | 14 | 4% | 100% |

### 2.5 Analysis of synonymous mutations in N ORF suggests implication of ZAP in progressive loss of CpG

We have then studied the nucleotidic patterns preceding, along the viral sequence, the CpG dinucleotides lost in Syn-SNV encountered so far. In N ORF, the ORF with largest density of CpG decreasing syn-SNV as shown in Table 3, 24 syn-SNV removing one CpG have been found. The nucleotide motifs preceding these loci are listed in the top 19 lines of Table 4 (for some loci, more than one syn-SNV removed the same CpG), together with their positions along N ORF of SARS-CoV-2 and their number of occurrences in the sequence data. 7 out of 19 of these loci, which represent 71% of total syn-SNV removing a CpG (1587 out of the 2239), correspond to a motif of the type CnxGxCG, where nx is a spacer of n nucleotides and were identified as ZAP binding patterns in [9]. The binding affinity of ZAP to the motifs was shown to depend on the spacer length, *n*, with top affinity for *n* = 7 [9] (see Table 4). Notice that 43% (3 out of the 7) of the CpG-suppression related motifs in SARS-CoV-2 correspond to *n* = 7. Other motifs of the type CnxGcCG are also present in SARS-CoV-2, but their CpG is not lost in sequence data, see last 5 lines of Table 4; the dissociation constants associated to their spacer lengths are on average larger than the ones of the motifs showing CpG loss.

**Table 4:**
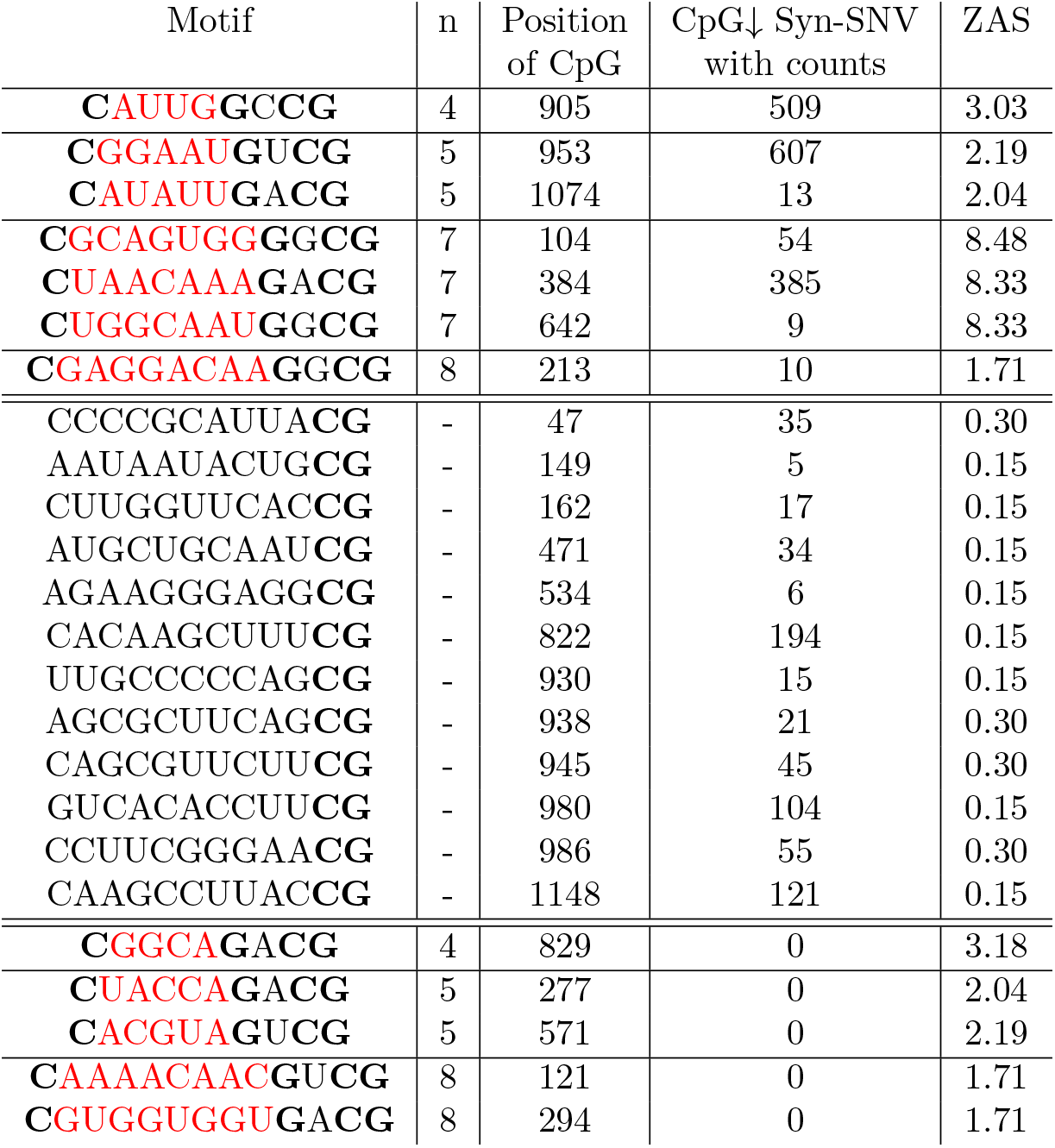
Analysis of nucleotidic motifs preceding CpG in the N ORF and their ZAP affinity score. The top 7 lines show subsequences of N ORF (of the Wuhan ancestral strain, GISAID ID: EPI ISL 406798) of the type CnxGxCG, where the spacer nx (highlighted in red) includes *n* = 4, 5, 7 or 8 nucleotides, for which the CpG dinucleotide was lost in one or more of the syn-SNV. These motifs were shown to be binding patterns for the ZAP protein in [9]; the dissociation constants were measured for repeated A spacers, with values (in *μ*M) *K_d_*(4) = 0.33 ± 0.05, *K_d_*(5) = 0.49 ± 0.10, *K_d_*(7) = 0.12 ± 0.04, *K_d_*(8) = 0.64 ± 0.14, [9]. The next 12 lines show the other CpG lost through mutations and their 10 preceding nucleotides, which do not correspond to motifs tested in [9]. The last 5 lines show other subsequences in the N protein corresponding to ZAP binding motifs [9], but for which no loss of CpG is observed in the sequence data. The column ZAS gives the score associated to the subsequence considered, computed from the above dissociation constants (see Methods for technical details). Data from GISAID [32], see Methods Sec. 4.6 for details on data analysis (last update 05 Oct 2020).

From the spacer-length dependent binding affinity given in [9] (see Table 4) we have computed a score, which we call ZAP affinity score (ZAS), which is related to the probability of having at least one ZAP bound to such motif (see Methods Sec. 4.4). The ZAS computed in sliding windows across the genome, is presented in Fig. 4a (top plot), from which it is apparent that N ORF is the richest region in motifs of the form CnxGxCG, with the largest ZAS. Our analysis is confirmed in Table 5, which reports all syn-SNV removing CpG following an extended sequence motif. Even if N ORF represents only 4% of the total sequence length, 18% of extended motifs CnxGxCG and 26% (58% with counts) of syn-SNV removing a CpG on an extended motif are on this region. In contrast, only 2 extended motifs of type CnxGxCG were found in 5’UTR even if many repeated CpG at short interspace were present, see Suppl. Table SI.1.

**Table 5:** ZAP affinity score and Syn-SNV removing CpG dinucleotides preceded by ZAP binding motifs across the SARS-CoV-2 genome. The table gives for each region with at least one motif of the form CnxGxCG with *n* = 4, 5, 6, 7 or 8 (CpGext) the length L, the number of CpGext and their number per unit length, the ZAS for unit length (ZAS/L), the number of syn-SNV removing a CpG preceding by an extended motif (CpGext↓SNV), and their fraction with respect the total number of CpG–decreasing syn-SNV. Additional information are given in Suppl. Table SI.2. Numbers and percentage of CpGext↓SNV are given with and without taking into account SNV counts. ORFs are sorted according to the density of CpGext removing SNV (CpGext↓SNV/L). On 5’ and 3’-UTRs there are no synonymous restriction on SNV.

|  | L | CpGext |  | ZAS/L | CpGext↓SNV |  |
| --- | --- | --- | --- | --- | --- | --- |
|  |  | num | /L |  | num | /CpG↓SNV |
| **5'-UTR** | 211 | 2 | 1.4% | 2% | 3 | 16% |
| with counts: |  |  |  |  | 88 | 0.2% |
| **N** | 1260 | 12 | 1.0% | 4% | 10 | 42% |
| with counts: |  |  |  |  | 1587 | 71% |
| **ORF7a** | 336 | 2 | 0.6% | 3% | 1 | 20% |
| with counts: |  |  |  |  | 17 | 7% |
| **ORF1a** | 13203 | 25 | 0.2% | 1% | 15 | 17% |
| with counts: |  |  |  |  | 726 | 18% |
| **ORF1b** | 8088 | 16 | 0.2% | 1% | 9 | 19% |
| with counts: |  |  |  |  | 317 | 4% |
| **S** | 3822 | 3 | 0.1% | 0.3% | 1 | 6% |
| with counts: |  |  |  |  | 25 | 4% |
| **ORF3a** | 828 | 4 | 0.5% | 2% | 0 | 0% |
| with counts: |  |  |  |  | 0 | 0% |
| **M** | 669 | 2 | 0.3% | 2% | 0 | 0% |
| with counts: |  |  |  |  | 0 | 0% |

N ORF and M ORF show a similar CpG force (see Table 3), but have a large difference in CnxGxCG-like motif content, as shown in Table 5. Remarkably, when taking counts under account, the number of CpG-decreasing mutations occurring in N ORF, out of which 71% are on CnxGxCG-like motifs, is 10-fold more than that occurring in M ORF. These results support the existence of early selection pressure to lower CnxGxCG-like motifs in N ORF, where they are particularly concentrated.

### 2.6 Model is able to discriminate observed and non-observed single nucleotide variants among early synonymous mutations

Our model can be further used to predict the odds of synonymous mutations, either implying CpG or not, from the ancestral SARS-CoV-2 (GISAID ID: EPI ISL 406798) sequence. For this purpose, we introduce a synonymous mutation score (SMS), defined in Methods Sec. 4.3, whose value expresses how likely a mutation is to appear under the joint actions of the CpG force, the codon bias (cb)^3^, and the transition-transversion bias (ttb)[36]^4^. For synonymous mutations that do not affect CpG the only mutational driving factors in our model are the codon bias, and the transition-transversion bias, which are global drives on the genome. Synonymous mutations changing CpG are also driven by the local force to increase or decrease CpG content depending on the CpG mutational drive in the region under consideration (see Fig. 4a and Table 3).

Figures 5a and 5b show SMS along, respectively, the N and S ORF, for all the observed syn-SNV lowering (blue), increasing (red), or leaving unchanged (gray) CpG content, along with their counts in the sequence sample. In N ORF as in S ORF the majority of syn-SNV have high SMS, validating the model predictions. The sign and amplitude of the SMS for CpG–affecting syn-SNV in Figs. 5a, 5b result from the interplay between the virus codon bias with the CpG drive in the model: the virus codon bias, computed on the whole genome which is globally low in CpG content, already favours synonymous mutations removing CpG. Due to the codon bias, both in N ORF and S ORF, syn-SNV removing CpG tend to have a positive SMS while syn-SNV adding CpG tend to have a negative SMS (see Suppl. Fig. SI.11 for SMS computed without CpG drive along N ORF and S ORF).

**Figure 5:**
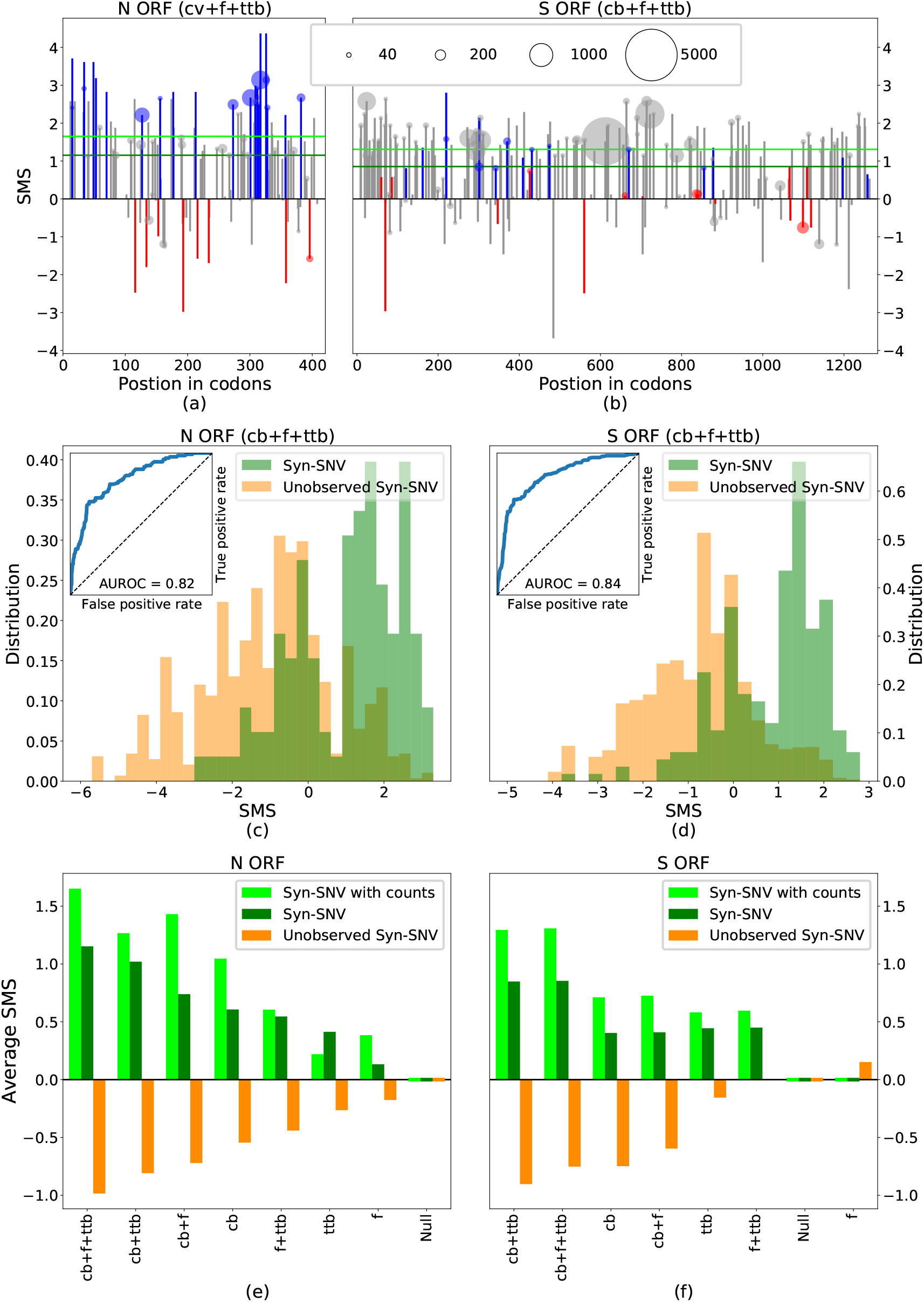
Synonymous Mutational Score (SMS) differentiates Syn-SNV from unobserved Syn-SNV. (a), (b): Synonymous Mutational Score (SMS) for Syn-SNV with the full model including codon bias (cb), CpG force (f) and transition-transversion bias (ttb) in the N and S ORFs. Blue, red, gray bars denote mutations decreasing, increasing or leaving unchanged the CpG content. The area of circles, shown on SNV observed more than 20 times in the dataset, is proportional to the SNV count. Green, horizontal lines are the average SMS of the Syn-SNV with (dark green) and without (light green) counts. (c), (d): histograms of SMS distribution for observed (green) and unobserved (orange) Syn-SNV in the N and S ORFs with the full model (cb+f+ttb). The corresponding ROC curve is given as an inset, together with the AUROC. (e), (f): Average SMS for Syn-SNV (dark green), Syn-SNV with SNV-counts (light green), and for unobserved Syn-SNV (orange), computed with the full model and all possible reduced models. In the null model (Null) all synonymous mutations are equiprobable. Models are ranked according to the difference of average SMS for observed and unobserved Syn-SNV. Data from GISAID [32], see Methods Sec. 4.6 for details on data analysis (last update 05 Oct 2020). Wuhan ancestral genome has GISAID ID: EPI ISL 406798. SNV with less than 5 counts are considered as unobserved Syn-SNV.

The large and negative CpG drive in N ORF adds to the codon bias trend. As shown in Fig. 5a, it further raises the SMS of CpG-removing syn-SNV, and further decreases the negative SMS of CpG-adding syn-SNV, in agreement with the data. The resulting SMS amplitude for CpG affecting mutations is generally larger than for mutations non affecting CpG. On the opposite, in S ORF, as shown in Fig. 5b (see also Suppl. Figs. SI.11c and SI.11d), the positive drive acts against the codon-bias trend, reducing or raising the SMS for, respectively, CpG-decreasing or -increasing syn-SNVs. In some loci, the CpG drives is strong enough to reverse the sign of the SMS for CpG-increasing syn-SMS, and the SMS become positive. The resulting SMS amplitude for CpG–affecting mutations is generally smaller than for mutations leaving CpG content unchanged, in agreement with the observation than CpG–affecting syn-SNV are rare in S ORF.

To make our arguments more quantitative, we tested the ability of our model to discriminate between observed and non–observed syn-SNV in sequence data collected so far. For the sake of clarity, non–observed syn-SNV refers to the set of possible synonymous mutations that have been not observed so far in the sequence data (or observed very rarely, i.e. with less than 5 counts). In Figs. 5c and 5d we show that the distribution of SMS for Syn-SNV is shifted to higher values compared to its counterpart for non observed Syn-SNV, both in N ORF and S ORF. Hence, our model is able to statistically discriminate between Syn-SNV and non observed Syn-SNV (ANOVA F-test: 1085 for N, 5590 for S). Moreover, when ranking all possible synonymous mutations in decreasing order according to their SMS and considering them as true predictions if they have been observed in sampled sequences so far, and thus correspond to a Syn-SNV, or false predictions if the have not been observed in the collected sequences, we obtained very good classification performances (AUROC=0.82 for N, 0.84 for S). As a complementary test we give in Suppl. Fig. SI.8 the positive predicted value (PPV) as a function of the number of predictions showing that about 85% of the top scored 90 and 150 possible mutations, in N ORF and S ORF respectively, are Syn-SNV.

To identify what ingredient in the model is responsible for correctly distinguishing between observed and non-observed synonymous mutations, we compare the discriminatory performances of the full model (including the codon bias, the transversion-transition bias and the CpG drive) with all model variations obtained removing one ingredient at the time, up to a null model in which all synonymous mutations are equally likely and all have zero SMS. In Figs. 5e and 5f we compare, for the different models, the average SMS for syn-SNV, obtained both with and without their counts in the sequence sample (see horizontal light and dark green lines in Fig. 5a and Fig. 5b), with the average SMS for the non observed syn-SNV. The models are ranked by the difference of average SMS between observed and non–observed syn-SNV. As expected the null model is unable to distinguish between the two sets, assigning vanishing average SMS to both. The gap between the average SMS of observed and non–observed syn-SNV progressively increases when introducing back the different ingredient of the model, up to the full one.

We have checked that the choice of the viral codon bias is important for such discriminatory ability. When using the human codon bias instead of the viral one the model based only on the human codon bias behaves similarly to the uniform bias model; the full model is again the best model but with a smaller difference in averages SMS (Suppl. Fig. SI.6). Classification performances are, on the contrary, unaffected when changing the equilibrium force to the values already discussed, such as the the global force SARS-CoV-2 or the average force computed on transcripts encoding human type-I IFNs (Supplementary Fig. SI.7). It is worth noticing that models based on viral codon and transition-transversion biases alone (without the force) already provide good discriminatory performances in terms of classification of SNV (AUROC and F-score). This result is expected from the predominance of Syn-SNV not affecting CpG occurences, especially in S ORF. In addition, as the viral sequence has low global CpG content, the viral codon bias favors synonymous mutations that lower CpG, in agreement with Figs. 5e and 5f.

Table 6 gives the average SMS and differences, as well as AUROC and ANOVA tests, for all ORFs and 5’ and 3’UTR. We consistently find that the full model is always a good model to describe syn-SNV observed in the early evolution of SARS-CoV-2. There is a clear separation between observed and unobserved syn-SNV average SMS with large AUROC for all the ORFs and UTRs (*≥* 0.68 for all the ORFs and UTRs and ≥ 0.71 for the regions with a better mutational statistics, with at least 20 syn-SNV) and a large ANOVA F value (≥ 3 for all the ORFs and UTRs and ≥ 13 among regions with at least 20 syn-SNV); the ORF1a, S ORF and ORF7b are the only regions for which the average difference between SMS of observed and non-observed syn-SNV computed with no CpG force is slightly larger. This again suggests that, for the S ORF, Syn-SNV are only marginally affecting CpG motifs and are mainly driven by codon bias and transversion-transition bias.

**Table 6:**
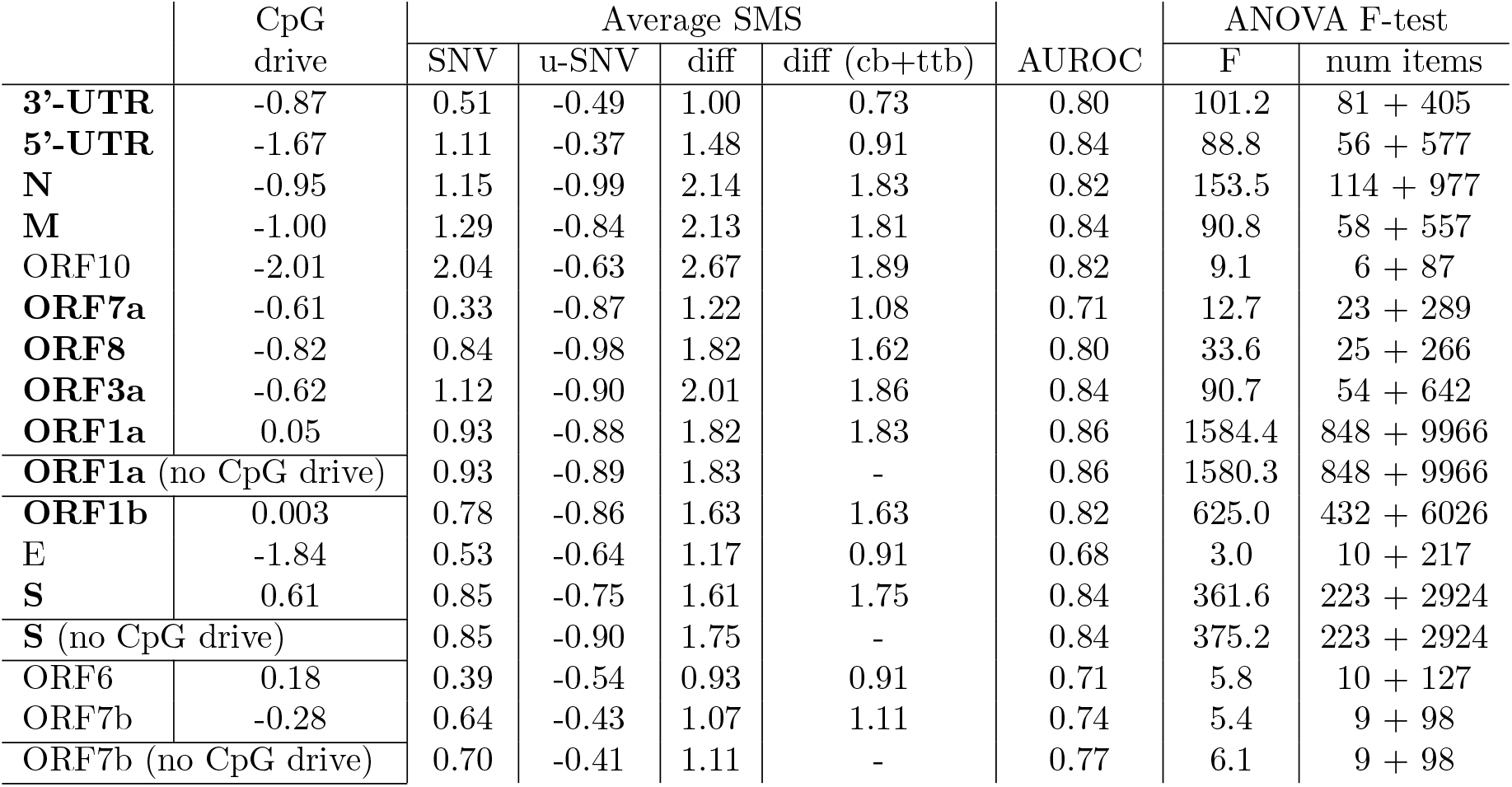
Model performance in predicting SNV on SARS-CoV-2 UTRs and ORFs. The table gives for the SARS-CoV-2 UTR and ORFs the CpG drive (*f_eq_* − *f*), the average Synonymous Mutational Score (SMS) for Syn-SNV and unobserved Syn-SNV (u-SNV) in the data collected so far (SNV with less than 5 counts are considered as unobserved), with the full model including codon bias (cb), CpG drive (f) and transition-transversion bias (ttb) in the N and S ORFs, the average SMS difference (diff) between the syn-SNV and the unobserved syn-SNV. To assess the role of the force in the model, we also provide difference in averaged SMS for the model not taking into account the CpG drive (diff (cb+ttb)). For ORF1a, S and ORF7b the presence of the CpG drive slightly decreases the average SMS difference, hence we provide the AUROC and ANOVA F-test results also with the model without CpG drive (see Suppl. Table SI.3 for all ORFs and UTRs). All ANOVA F values are significant (p-value < 0.05) at exception for E ORF (p-value=0.06). The number of items (num items) in the two sets to compute F are the Syn-SNV + all the possible unobserved Syn-SNV. SNV on 5’ and 3’-UTRs have not the synonymous restriction.

Remarkably, the ranking of models and their discriminatory performances are essentially the same when taking into account or not the counts of the Syn-SNV, even if a net increase in the average Syn-SNV SMS score is present for models with the CpG drive in the N ORF, when counts are considered. On the one hand, ignoring counts leads to conservative estimates of the SMS, as mutations which are fixing in the population are not weighted more than less frequent mutations. On the other hand, SMS based on sequence counts better discriminate observed and unobserved Syn-SNV in most of ORFs and UTRsAs (Supp. Table SI.4), but are likely plagued with phylogenetic and sampling biases. We expect that SMS, when deriving mutational fitness from phylogeny [37] and correcting for sampling bias, will lie in between the two limit-case SMS discussed above. Finally, we checked the consistency of our results at different times since our first analysis (dated 05 May 2020, see Suppl. Fig. SI.10).

## 3 Discussion

The present work reports analysis of dinucleotide motif usage, particularly CpG, in the early evolution of SARS-CoV-2 genomes up to October 2020. First, a comparative analysis with other genomes shows that the overall CpG force, and the associated CpG content are not as large as for highly pathogenic viruses in humans (such as H1N1, H5N1, Ebola and SARS and MERS in the *Coronaviridae* family). However, the CpG force, when computed locally, displays large fluctuations along the genome. This strong heterogeneity is compatible with viral recombination, in agreement with the hypothesis stated in [38]. The degree to which this heterogeneity in any way reflects zoonotic origins should be further worked out using phylogenetic analysis. In particular, the segment coding for the Spike protein has a much lower CpG force. The S protein has to bind ACE2 human receptors and TMPRSS2 [18, 34]. A fascinating reason that could explain the low CpG force on this coding region is that it may come (at least in part) from other coronaviruses that better bind human entry receptors [38, 39]. Other regions, in particular the initial and final part of the genome including the 5’ and 3’ UTR and N ORF, are characterized by a larger density of CpG motifs (and corresponding CpG force), which are comparable to what is found in SARS and MERS viruses in the *Betacoronavirus* genus. Interestingly the initial and final part of the genome are implied in the full-genome and subgenomic viral replication. In particular, the coding region of the N protein and its RNA sequence, present in the 3’UTRs of all SARS-CoV-2 subgenomic RNAs, has been shown in [28] to be the most abundant transcript in the cytoplasm. The high concentration of N transcripts in the cytoplasm could contribute to a dysregulated innate immune response. A mechanism generating different densities of PAMPs being presented to the immune system at different points in the viral life cycle can affect immune recognition and regulation. The precise way this can contribute to immuno-pathologies associated with COVID-19 and how this is related to the cytokine signaling dysfunction associated with severe cases [17], needs further experimental investigation.

The analysis of the evolution of synonymous mutations since the outbreak of COVID-19 shows that mutations lowering the number of CpG have taken place in regions with higher CpG content, at the 5’ and 3’ ends of the sequence, and in particular in the N protein coding region. The sequence motifs preceding the loci of the CpG removed by mutations match, especially in N ORF, some of the strongly binding patterns of the ZAP protein [9]. Natural sequence evolution seems to be compatible for protein N with our model, in which synonymous mutations are driven by the virus codon bias and the CpG forces leading to a progressive loss in CpG. These losses are expected to lower the CpG forces, until they reach their equilibrium values in human host, as is seen in coronaviruses commonly circulating in human population [40]. More data, collected at an unprecedented pace [31, 32, 41], and on a longer evolutionary time are needed to confirm these hypothesis. Since the data collected are likely affected by relevant sampling biases, a more precise analysis of synonymous mutations could be carried out using the available phylogenetic reconstruction of viral evolution [42]. Nevertheless our results seem robust, since they are consistent both considering unique mutations and all collected synonymous variants. They coherently point to the presence of putative mutational hotspots in the viral evolution. While the results presented here are preliminary due to the early genomics of this emerging virus, they have been confirmed by incoming sequence data since our first analysis (dated 05 May 2020, see Suppl. Fig. SI.10) and they point to interesting future directions to identifying the drivers of SARS-CoV-2 evolution and building better antiviral therapies. In this work we focused on synonymous mutations, but it would interesting to extend our fixed amino-acid model for viral evolution to take into account non-synonymous mutations and to further model transmission and mutations (in the presence of a proofreading mechanism [43]) processes in SARS-CoV-2 to predict the time scale at which natural evolution driven by host mimicry would bring the virus to an equilibrium with its host [4, 5].

After our work was posted on the bioRxiv, R. Nchioua and colleagues have shown the importance of ZAP in controlling the response against SARS-CoV-2 [44] by demonstrating that a knock-out of this protein increases SARS-CoV-2 replication. The interaction between SARS-CoV-2 and ZAP has also been observed with unbiased methods in another recent work [45]. This finding supports our prediction that recognition of SARS-CoV-2 by ZAP imposes a significant fitness cost on the virus, as demonstrated by its early evolution to remove ZAP recognition motifs. Two other recent theoretical works [46, 47], corroborate our results showing that at the single nucleotide level there is a net prevalence of C→U synonymous mutations (the most common nucleotide mutation which may cause a CpG loss) in the early evolution of SARS-CoV-2. Moreover a recent analysis of the immune profile of patients with moderate and severe disease revealed an association between early, elevated cytokines and worse disease outcomes identifying a maladapted immune response profile associated with severe COVID-19 outcome [48].

## Acknowledgments

We thank Nicolas Vabret for discussions and reading of the manuscript, Eddie Holmes and Marta Luksza for helpful exchanges. We gratefully acknowledge the authors, originating and submitting laboratories of the sequences from GISAIDs EpiCoV(TM) Database on which this research is based. This work was partially supported by the ANR-19 Decrypted CE30-0021-01 grant and by the Fondation de la Recherche Médicale (FRM) (ANR-Flash Covid, project SARS-Cov-2immunRNAs). B.G. was supported by National Institutes of Health grants 7R01AI081848-04, 1R01CA240924-01, a Stand Up to Cancer Lustgarten Foundation Convergence Dream Team Grant, and The Pershing Square Sohn Prize Mark Foundation Fellow supported by funding from The Mark Foundation for Cancer Research.

## 4 Material and Methods

### 4.1 CpG density versus local and global forces

Throughout this work we used CpG forces to quantify the CpG content of a given sequence. Here we want to compare this approach with the simple count of CpG motifs in the sequence. In Suppl. Fig. SI.2 we show that some of our results, such as the large fluctuations of the CpG content across the SARS-CoV-2 genome, are also apparent from a simple motif density analysis. However, this counting strategy is not suitable to make comparisons among viruses of different families, mainly because of the different lengths and usage biases of viral genomes. Moreover, without the statistical framework at the basis of the CpG force, it is very difficult to take into account the many constraints acting on a genetic sequence, notably the constraint on the amino acids that have to be coded for in the sequence. The force formalism is, therefore, much more flexible and allows us to introduce in a theoretically grounded way the synonymous mutation score (see Sec. 4.3) which we used to characterize mutations which are likely to happen. The drawback of such a formalism is the quite large number of extra choices that have to be done, and which can influence the result. These choices are discussed in the following section.

### 4.2 Force computation

The technique at the core of many of the analyses made here is taken from [4]. Here we briefly review this technique, starting from its non-coding version which takes as reference bias the nucleotide bias and then describing the coding version which take as reference bias the codon bias at fixed amino-acid sequence.

#### Force computation in non-coding case

Given a motif *m* and a sequence *s*_0_ = {*s*_1_, …, *s_N_* } of length *N*, we consider the ensemble of all sequences with length *N*, which we denote with 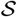, and we suppose the probability of observing *s* out of this ensemble to be

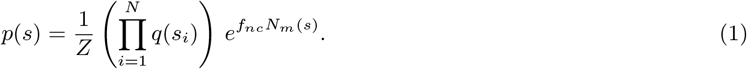

Here, *q*(*s_i_*) is the nucleotide bias, that is the probability of the i-th nucleotide being *s_i_* (for example, we always used in this work the human frequency of nucleotides as *q*(*s_i_*)), *f_nc_* is the force we want to compute (the subscript *nc* stands for non-coding), and *N_m_* is the number of times the motif *m* appears in the sequence. *Z* is the normalization constant, that is

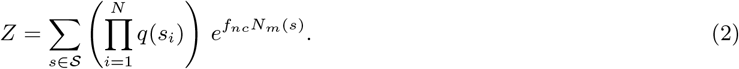

Therefore the force *f_nc_* is a parameter which quantifies the lack (if negative) or abundance (if positive) of occurrences of *m* with respect to the number of occurrences due to the local probabilities *q*(*s_i_*). We can fix *f_nc_* by requiring that the number of motifs in the observed sequence, *N_m_*(*s*_0_) = *n*_0_, is equal to the average number of motifs computed with the probability Eq. 1, ⟨*n*⟩, that is

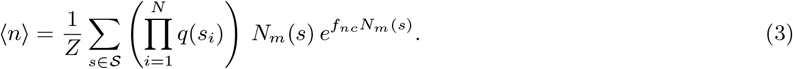

Notice that this is equivalent to the request that *f_nc_* is so that probability of having observed *s*_0_ is maximum.

Let us focus now on the specific case of a dinucleotide motif, that is our motif *m* consists of the pair *ab*, where *a* and *b* are two consecutive nucleotides (for example, *a* = *C* and *b* = *G* for the CpG motif). In this case, within an approximation discussed in the SI, Suppl. Sec. SI.3, the force computed as above turns out to be the logarithm of the relative abundance index, that is

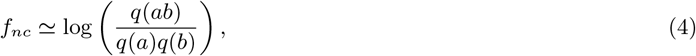

where *q*(*ab*) is the number of motifs *ab* divided by the total length of the sequence *N* . In Suppl. Fig. SI.12 we tested the accuracy of this approximation in our specific case. As it is clear from Eq. (4), the choice of the nucleotide bias *q*(*C*) affects the absolute value of the forces but not the difference between forces computed on different viral genomes using the same reference bias. We have chosen as reference nucleotide bias the human nucleotide bias (computed on all the genome, or on the coding DNA only). This choice can be then replaced by any other reference bias (possible choices include the codon bias computed on the ancestral SARS-CoV-2 sequence, or other human Coronavirus viral sequences or the one computed on RNA transcripts of human type-I IFNs, at the core of of innate immune response) and will shift the values of the forces without affecting the ranking of the force on different viral sequences, see Fig. 2 and Table 1.

#### Force computation in the coding case

Our technique can be generalized to coding sequences at fixed amino acid sequence and codon bias. In this case, we write each sequence *s* as a series of codons, and its probability is defined as

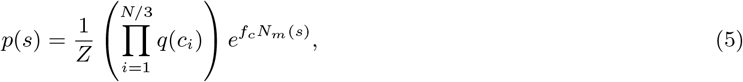

where now the bias takes the form of a codon usage bias, and the normalization constant *Z* changes accordingly into a sum over all possible synonymous sequences. The subscript *c* stands for coding, and differentiates this force from the non-coding one. The force *f_c_* can be computed, analogously with the procedure for the simpler case, by requiring that the number of motifs observed in *s*_0_ is equal to the statistical average performed with Eq. 5, as described in detail in [4]. As shown in Fig. 3a, the CpG force at fixed amino acids are roughly comparable to the one at fixed nucleotide bias when computing the nucleotide bias on human coding sequences.

The force computed in the coding (or non-coding case) is an useful tool to determine the content of a given dinucleotide, while taking into account a number of constraints.

### 4.3 Definition of synonymous mutation score

We use the ideas introduced above in Sec. 4.2 to introduce a model in order to assign a score, which we call synonymous mutation score (SMS), to each possible single-codon synonymous mutation of an ancestral sequence. We consider a system evolving for a small time scale, and a mutation which changes the *i*-th codon *c_i_* into a synonymous 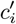. The transition probability, that is the probability of observing the mutation, for such evolution can be decomposed in the product of two evolution operators: The first 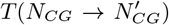 representing the change in the number of CpG motifs in the mutated sequence, and the second 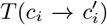 representing the gain in mutating the particular codon in position *i*.

The first operator can be computed from the dynamical equation introduced in [4] for the evolution of the CpG number *NCG* of a sequence under a initial force^5^ *f* through a equilibrium force *f_eq_* :

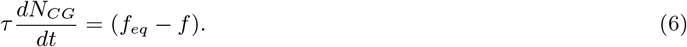

The equilibrium force can be computed on a viral strain which is supposed to be to the equilibrium with the human innate immune system, because it has evolved in the human host since a long time. Eq. 6 was used in [4] to describe the evolution of the CpG numpber in H1N1, taking as the equilibrium force the one of the Influenza B strain. In analogy with this approach we take here as *f_eq_* the average force calculated on coding regions of the seasonal hCoVs (that is hCoV-229E, hCoV-NL63, hCoV-HKU1, hCoV-OC43). Other possible choices are discussed in Sec. 4.5 (see also Suppl. Fig. SI.7). *τ* is a parameter determining the characteristic time scale for synonymous mutations. Based on Eq. (6) we define the transition operator for CpG number as

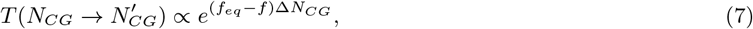

where 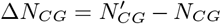. Notice that for all the synonymous mutations leaving unchanged the CpG number the above operator is one. The codon mutational operator is

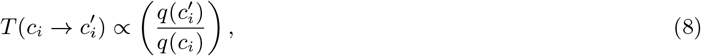

where *q*(*c_i_*) is the frequency of codon *c_i_* from the chosen codon usage bias. Putting together these two terms allows us to estimate how likely a specific synonymous mutation is to happen. The synonymous mutation score (SMS) accompanying a mutation is defined as the logarithm of this quantity,

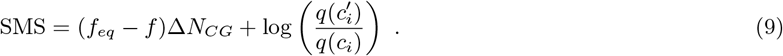

To conclude, we remark that different models can be used in the SMS computation, where a model is specified by giving the choice of including or not the force term, the choice of the equilibrium force to be used, the choice of including or not the codon bias term, and choice of the reference codon bias to be used.

#### 4.3.1 Adding transition-transversion bias to SMS

It is well known that transversions (i. e. mutations of purines in pyrimidines and vice-versa) are suppressed with respect to transitions (i. e. mutations of purines in purines or pyrimidines in pyrimidines).

We introduce here a simple way to account for transition-transversion bias in the model used to assign the SMS. We suppose that a mutation with *n* transversions happens 4 times less than a mutation with *n* − 1 transversions. This probability ratio, which is a standard value in the literature [36], has been recently shown to be close to the observed value for SARS-CoV-2 [49]. To include that in our model, consider mutating a codon *c* to *c*′, one of its synonymous codons. Let us suppose that the SMS for this event, computed with a given model, is SMS(*c, c*′). We then count the number of transitions, *n_trn_*, and the number of transversions, *n_trv_*, and modify the SMS into SMS’, so that

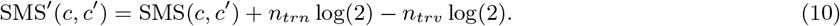

This choice is motivated by mainly two considerations: (i) in this way, a dynamical model where mutation probabilities are proportional to the exponential of SMS (as the one considered in Sec. 4.3 to justify the SMS itself) correctly gives a 4-fold probability to a transition than a tranversion (if the two mutations have the same SMS without this new term); (ii) the splitting on the extra term in a positive weight for transitions and a negative weight for transversions ensures that the average SMS before and after adding this term is comparable.

### 4.4 ZAP Affinity Score

We introduce the ZAP Affinity Score (ZAS) to roughly quantify a priori the likelihood of ZAP biding to a given region of RNA. ZAS is based on the dissociation constants obtained in vitro in [9]. Let us consider the case of a single motif (be it CG, or CnxGxCG, with n=4, 5, 6, 7 or 8), 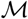, with dissociation constant *K_d_*. The association constant is then defined as

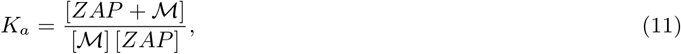

where 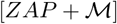 is the concentration of complexes, [ZAP] and 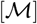 are the concentration of free molecules. Let us denote by [*ZAP*]_0_ and 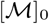 the total concentration of molecules (bound and unbound). If we suppose that only a small part of the available molecules form a complex, that is more specifically that *K_a_*[*ZAP*]_0_ « 1 and 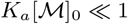, then *K_a_*[*ZAP*]_0_≃*k_a_*[*ZAP*] is the probability of binding. If we have *n* sites with association constants 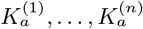, the probability of observing at least one ZAP bound to the RNA is

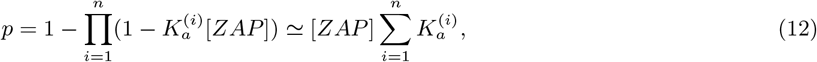

where we also used that *n* is sufficiently small so that 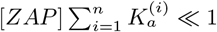. Finally, ZAS is defined as

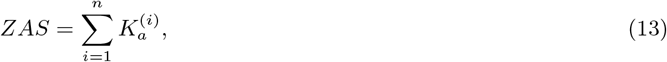

that is *p*/[*ZAP*]. While ZAS itself does not depend on [*ZAP*], its interpretation (and in particular its connection with the probability of binding) does, as it requires 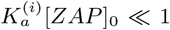, 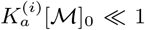, and 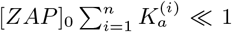. The 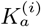 used here range from about 10^5^ (for the simple CpG motif) to 10^7^ (for C7xGxCG) *mol/L*. It is more difficult to estimate 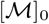 and [*ZAP*]_0_ during the infection. However, we hypothesize that these requirements are fulfilled in cells, and that our interpretation in terms of binding probability is acceptable.

### 4.5 Robustness of analysis with respect to choice of parameters

We discuss here how force values and SMS scores change by changing model parameters.

Parameters affecting the force values:

i. *Nucleotide, codon bias choices:* the most relevant effect due to this choice is a global shift of the force, as we show in Fig. 3a for the non-coding case, which does not change the ranking of forces when comparing different sequences using the same reference bias.
ii. *Choice of the length of the segment to compute the force*: the force is an intensive parameter. However, here we use the force to quantify the content of CpG motifs, which are quite rare. For this reasons, computing forces on small segments can lead to large negative values (the force is *−∞* when no CpG motif is present), and to unnatural fluctuations. For this reasons, to compute local force we fix large sliding windows of 3 kb, and we use Gaussian sliding averages to smooth the resulting curves. The effect of Gaussian smoothing and changing sliding window on the force are presented in Fig. SI.9.

Parameters affecting the SMS:

i. *The codon bias (or nucleotide bias for mutations in 5’ and 3’-UTRs)*: it is both present as a reference bias in the force computations for the CpG drive term, and directly used as a more generic evolutionary driver for the synonymous mutations. For the computation of forces in the CpG drive we have used the human codon or nucleotide bias as reference usage, but such choice is actually irrelevant because the drive is a difference of the segment force and the equilibrium one. The choice of bias is, on the opposite, very relevant for the choice of the synonymous mutations driver. Indeed, in Figs. 5e and 5f it is apparent that the virus codon bias alone gives to the model a certain ability to discriminate between observed an unobserved syn-SNV. We tested also the human codon bias which gives bad performances (see Suppl. Fig. SI.6).
ii. *Choice of equilibrium force*: this choice is arbitrary to a certain degree. We use as equilibrium force (computed with the human codon bias) −1.79 which is the average coding force hCoVs (229E, NL63, HKU1, OC43), since these viruses are well adapted to the human environment so likely a good equilibrium point for SARS-CoV-2. To check the effects of other choices of equilibrium forces, in Fig. SI.7 we performed the same analysis shown in Figs. 5e and 5f with other two possible choices of equilibrium forces: the global force of SARS-CoV-2 (−1.71, which is quite low, but slightly higher than the average of the seasonal hCoVs), and the average global force of INF-I transcripts (−2.89 which is much lower than that of seasonal hCoVs), see also Tables 1 and 2. While the SMS assigned to the mutations are in general different, especially when taking into account the counts in syn-SNV, and so the average SMS in figures, the ranking of the various model in term of average SMS difference (syn-SNV versus unobserved syn-SNV) is quite robust.
iii. *Presence of transition-transversion bias*: we observe that the presence of this term always increases the difference between the average SMS in observed and unobserved synonymous SNV. Two choices are needed to fix this term: the value the probability ratio of a transition with respect to a transversion (here we considered this ratio to be 4), and the specific way of realizing this bias by adding a bonus/penalty term to transitions and trasnversions. The latter choice is almost irrelevant when considering the differences of average SMS between observed and unobserved SNV.

### 4.6 Data Analysis

SARS-CoV-2 sequences are taken from GISAID [32]. We collected each sequence present in the database on 05 Oct 2020 (the most recent sequence was collected on 29 Sept 2020). Before any of our analyses, we discarded all the sequences where one or more nucleotides were wrongly read (other characters than A, C, G, T, U). This left us with 56045 SARS-CoV-2 sequences. To obtain Fig. 2 we considered, in addition to the SARS-CoV-2 sequences are taken from GISAID, other *Alphacoronavirus* and *Betacoronavirus* sequences (whole genomes and genes) which have been obtained from VIPR [31]. The pre-processing consisted again of discarding all the sequences with one error or more. After this process we collected 341 SARS, 48 MERS, 20 hCoV-229E, 48 hCoV-NL63, 14 hCoV-HKU1, 124 hCoV-NL63, 166 bat-CoVs and 5 pangolin-CoVs whole genomes. For Fig. 2b we used the largest number possible of sequences, up to a maximum of 100. For Fig. 2a (viral sequences) and Fig. 2c we chose a single sequence for each species. However, we checked that the result is qualitatively the same if we use other sequences from the same species for human coronaviruses. The curves in Fig. 2c are smoothed through a Gaussian sliding average (on windows of 3 kb, the Gaussian being centered in the window, normalized, with a standard deviation of 300 b). The ancestral SARS-CoV-2 sequence used throughout the work has been collected on 26-12-2019 (ID: EPI ISL 406798).

In Fig. 3a the SARS-CoV-2 sequence has been processed to ensure the correct reading frame. Therefore the ORF1ab gene is read in the standard frame up to the ribosomal shifting site, and it is read in the shifted frame from that site up to the end of the polyprotein. Moreover, the small non-coding parts between successive proteins have been cut, resulting in a loss of 634 nucleotides (including the 5’-UTR and 3’-UTR). A Gaussian smoothing has been performed to obtain the plotted CpG forces (as in Fig. 2c). To produce the bar plots in Figs. 3c and 3b we collected genes data on VIPR. Then we discarded as usual all the sequences with one or more errors, and we computed for each gene an average of up to 20 different sequences (coming from the same species). For some structural proteins we did not find 20 different genes but in any case the standard deviation of the averages of Figs. 3c and 3b is smaller than 0.02 (and, for most of the viruses, much smaller). In particular, we used 20 sequences of SARS-CoV-2, MERS, hCoV-NL63, hCoV-OC43 proteins, 14 sequences for hCoV-229E, 13 for hCoV-HKU1 and 4 for SARS. More detailed information about the genomes used in this work are given in Suppl. Sec. SI.1.

The mutations used for Figs. 4 and 5, have been collected by extracting ORFs from the SARS-CoV-2 sequence dataset, and comparing them with the Wuhan ancestral strain. ORFs with mutations too close to the start or end codon are not considered, together with ORFs with insertion/deletion, this filtering procedure leaving us with 48511 sequences to obtain mutation data. Mutations with less than 5 counts in different sequences are discarded. All curves in Fig. 4a are smoothed with the same Gaussian average used in Figs. 2 and 3. Finally, to get the mutation data in 5’UTR and 3’UTR we considered the UTRs of the Wuhan ancestral strain, and we compared them with those of other sequences. The number of nucleotides of the Wuhan ancestral considered part of 5’UTR and 3’UTR for the search for mutations in other sequences is given in Table 3. This length is chose so that a large number of uploaded sequences (about 50000) have a UTR of the same length or longer. In the UTR analysis, all observed mutations are considered “synonymous”. The code used to compute coding and non-coding forces is publicly available at https://github.com/adigioacchino/dinucleotide_forces.

## Supplementary Information

### SI.1 Genomes analyzed

Here we report some additional information about the genomes used in this work. The SARS-CoV-2 sequence used in Fig. 2 has GISAID accession ID: EPI ISL 420793. The other GenBank accession numbers for the specific genomes used in Figs. 2 are: AY427439 (SARS), NC 038294 (MERS), MF542265 (hCoV-229E), JX524171 (hCoV-NL63), KT779555 (hCoV-HKU1) and KF923918 (hCoV-OC43). For these figures, we choose the bat and pangolin sequences closest to the SARS-CoV-2 points in Fig. 2b (these two sequences are also known to be very similar to the SARS-CoV-2 genome from previous works [38]). These sequences have GISAID accession IDs EPI ISL 402131 (bat coronavirus sequence known with the name RaTG13) and EPI ISL 410721 (pangolin coronavirus sequence collected in 2019 in Guangdong).

The SARS-CoV-2 ancestral sequence, which has been collected in Wuhan on 26-12-2019, has GISAID accession ID: EPI ISL 406798. This sequence has been used as reference in Figs. 3, 4 and 5. For Figs. 3b and 3c we used specific sequences, with the following GenBank accession numbers: MT300186:28249-29508 (SARS-CoV-2), AY291315:28120-29388 (SARS), NC 038294:28565-29800 (MERS) and KT779555:28281-29606 (hCoV-HKU1) for the N protein; MT300186:21538-25359 (SARS-CoV-2), AY291315:21492-25259 (SARS), NC 038294:21455-25516 (MERS) and KT779555:22903-26973 (hCoV-HKU1) for the S protein.

### SI.2 Supplementary Figures

**Figure SI.1:**
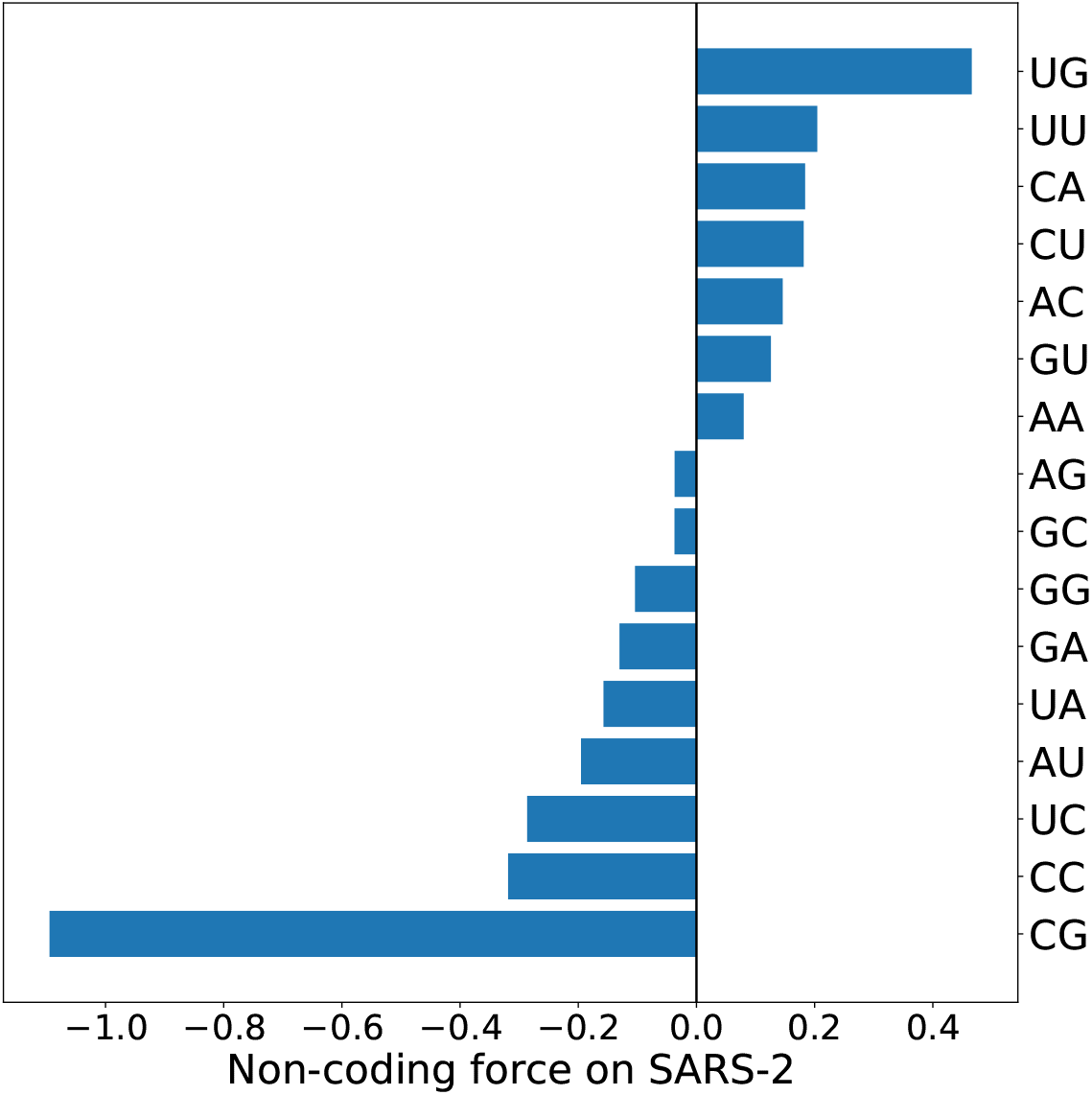
All dinucleotide non-coding forces computed on the whole SARS-CoV-2 genome. The CpG motif is the one with the largest non-coding force in absolute value, and the second one is UpG, which is one transition away from CpG.

**Figure SI.2:**
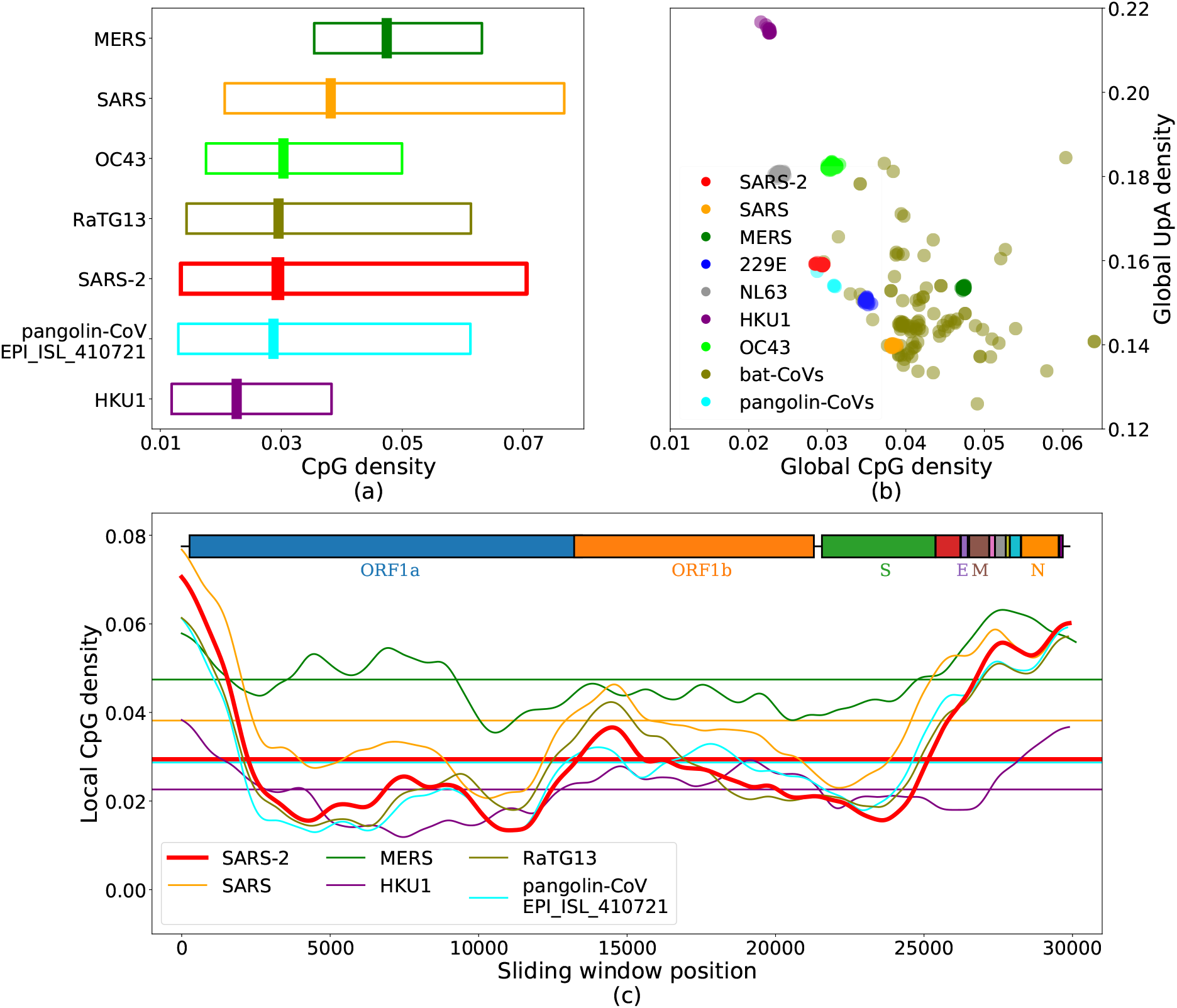
The same analysis performed in Fig. 2, but here we used CpG densities (CpG dinucleotides divided by the total number of dinucleotides), instead of CpG forces. The results obtained are qualitatively similar.

**Figure SI.3:**
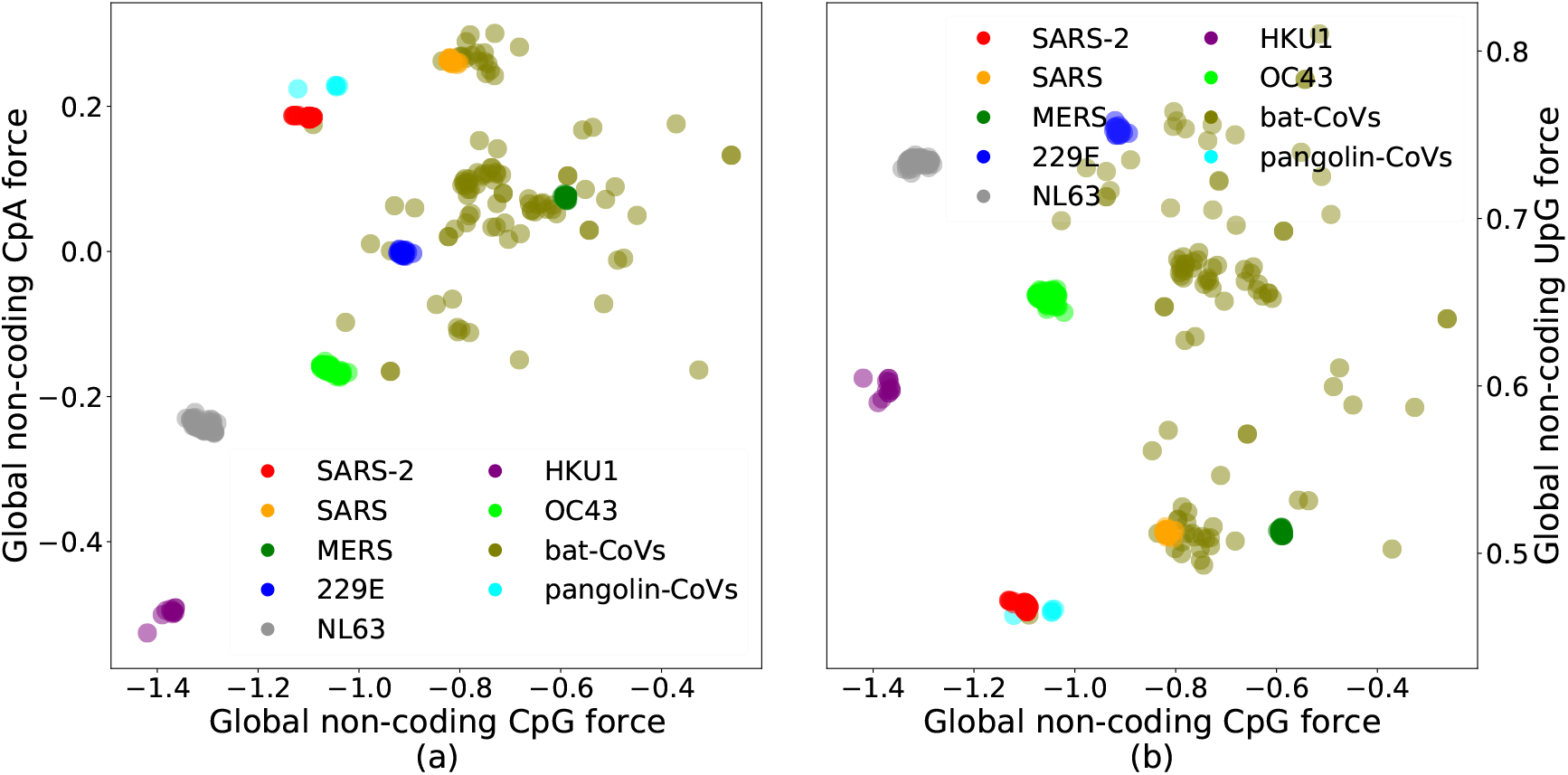
CpG versus CpA (left) and UpG (right) non-coding forces. Differently from the CpG versus UpA case (Fig. 2b), no clear correlations (*r*^2^ equal to 0.2 and 0.01, respectively, for the left and right plots) are found.

**Figure SI.4:**
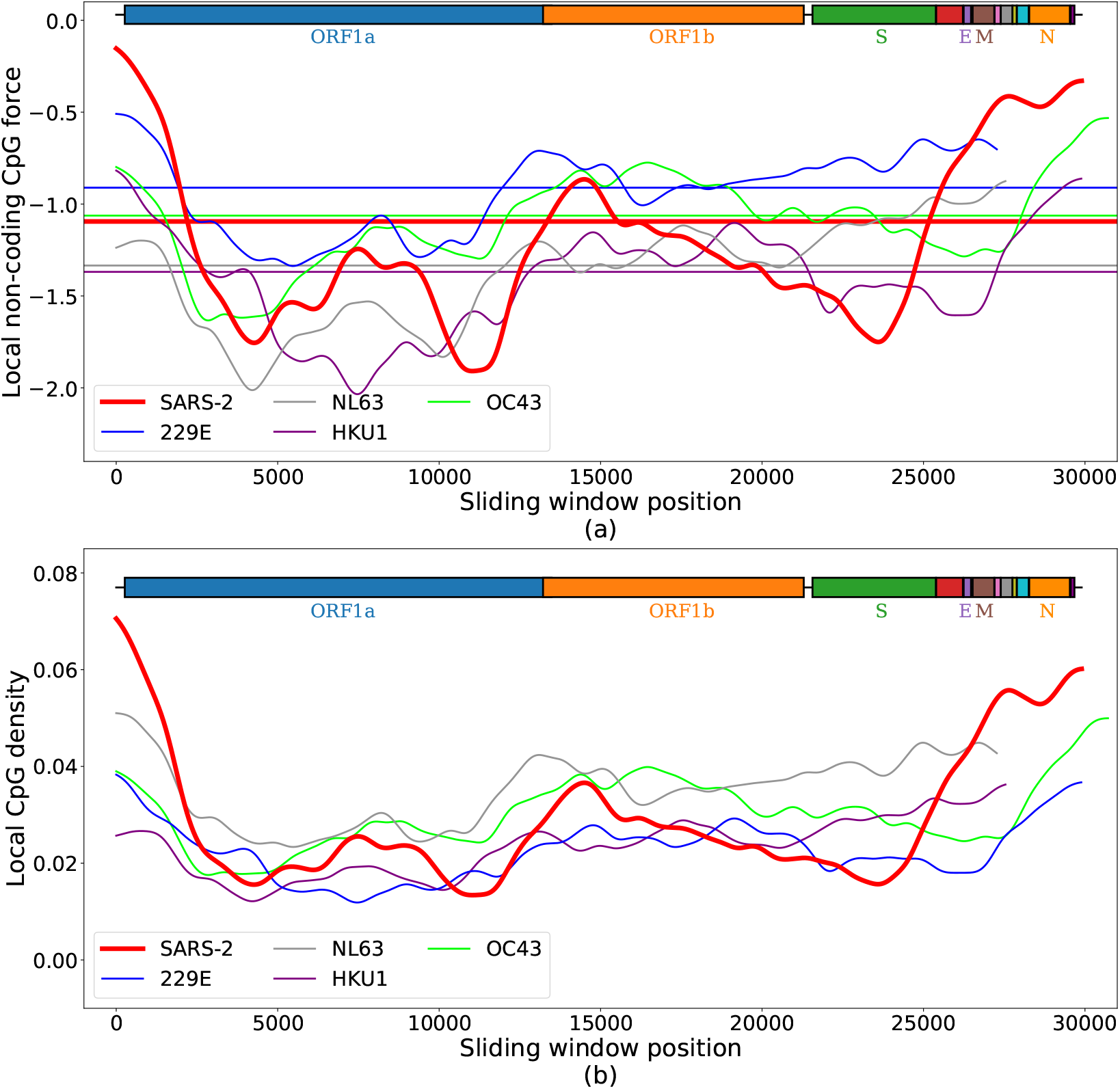
Supplement to Fig. 2 and Suppl. Fig. SI.2, where all the coronaviruses associated with circulating human strains are compared with SARS-CoV-2 in terms of CpG non-coding force (panel (a)) or number in fixed-length windows (panel (b)). Again, even though the final regions of the hCoVs has relatively high CpG force with respect to the other parts of their sequences, SARS-CoV-2 has a 3’ CpG force peak well above the final region of hCoV virus.

**Figure SI.5:**
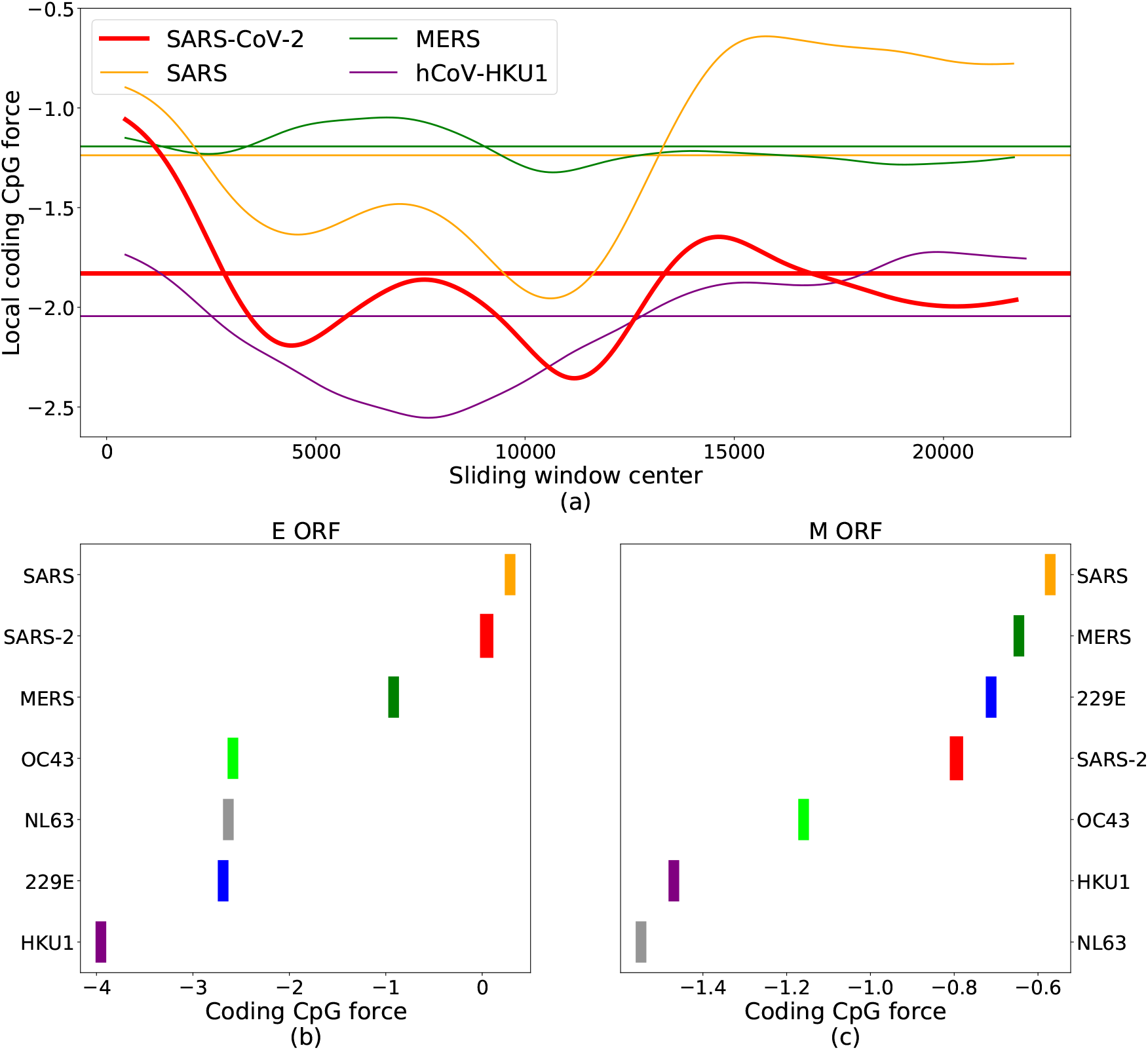
Extension of the comparison performed in Fig. 3. In panel (a) the CpG coding forces computed on the genome coding for polyprotein ORF1ab is compared among several coronaviruses and in panels (b) and (c) the structural proteins E (envelope) and M (membrane) are considered. Notice that, due to the small size of proteins E and M, only one window is used so the boxes collapse in one line corresponding to the global CpG force on the protein, computed as an average over 4 to 20 viral genomes.

**Figure SI.6:**
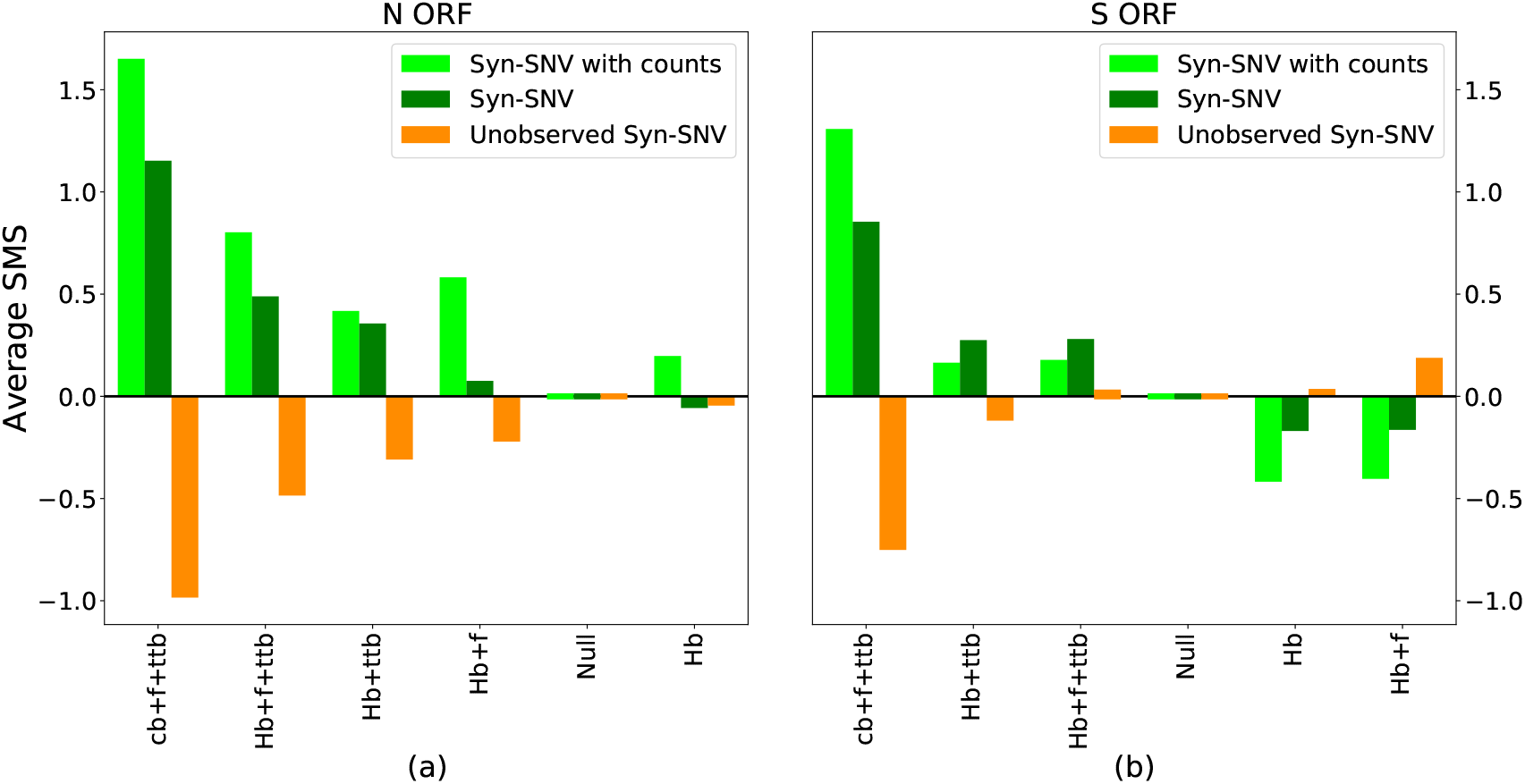
Supplementary information for Figs. 5e and 5f, where the model with human codon bias (Hb), with or without force and transition-transversion bias, is compared to the null model and the the model with virus codon bias (cb), force and transition-transversion bias, in terms of average SMS assigned to synonymous SNV in N and S ORFs.

**Figure SI.7:**
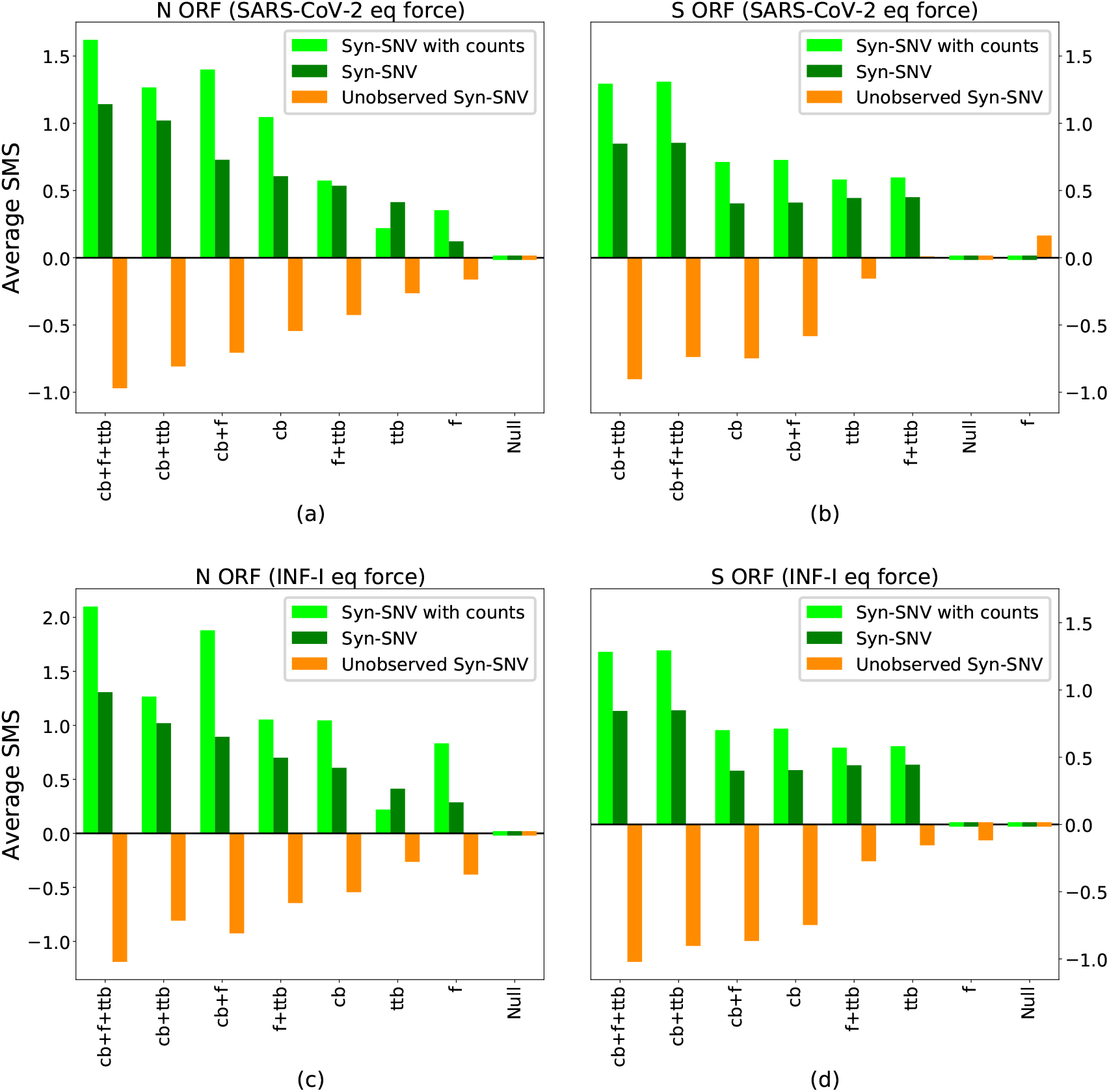
Same of Figs. 5e and 5f, for different choices of equilibrium force. (a), (b): the equilibrium force chosen is −1.71, that is the global coding force on SARS-CoV-2 (computed with the human codon bias as background). (c), (d): the equilibrium force chosen is −2.89, that is the average global non-coding force of human type-I IFNs (computed with the coding human nucleotides as background).

**Figure SI.8:**
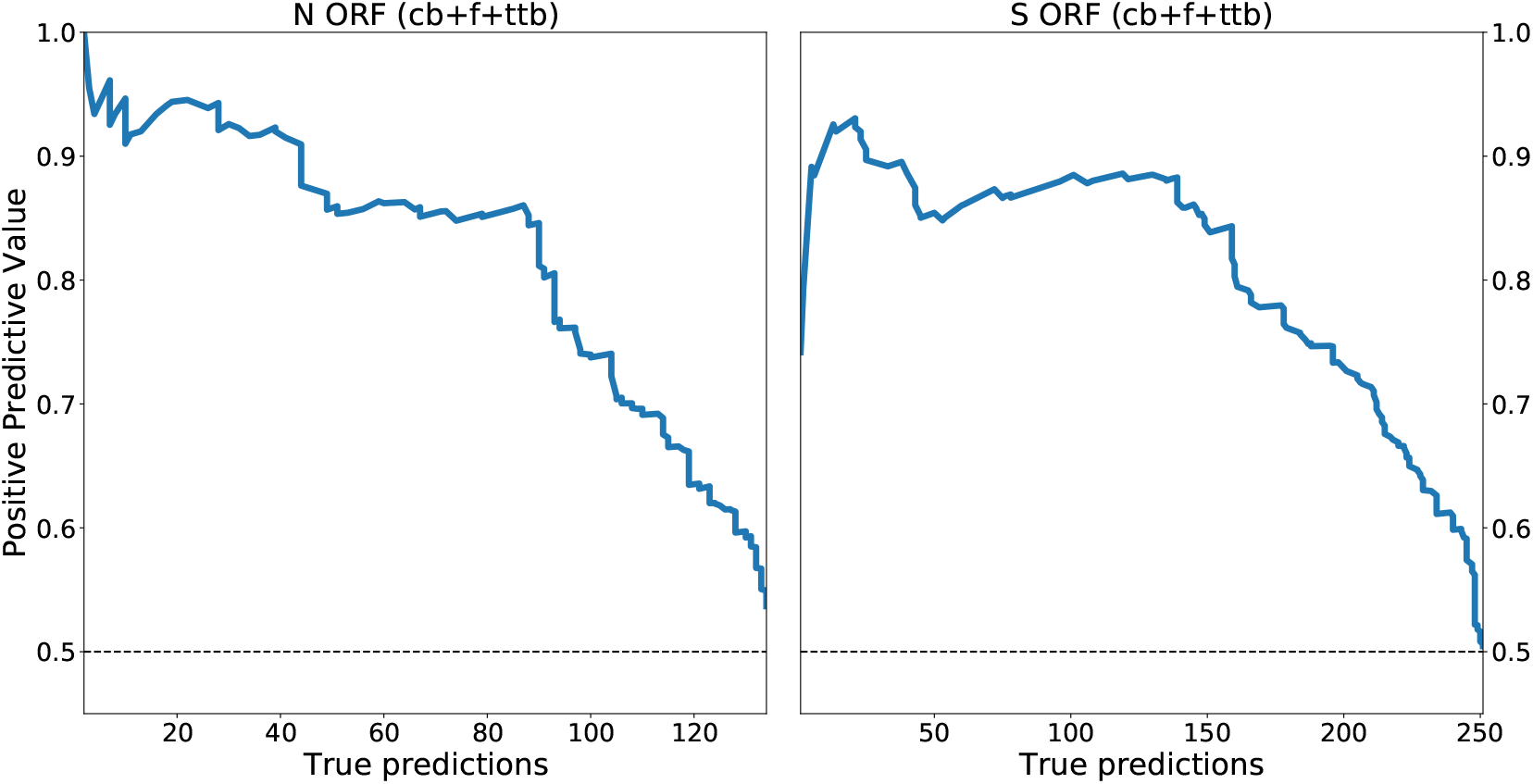
Positive Predictive Value for the model with virus codon bias, CpG force and transition-transversion bias, for N and S ORFs, as a function of the number of correctly predicted synonymous SNV.

**Figure SI.9:**
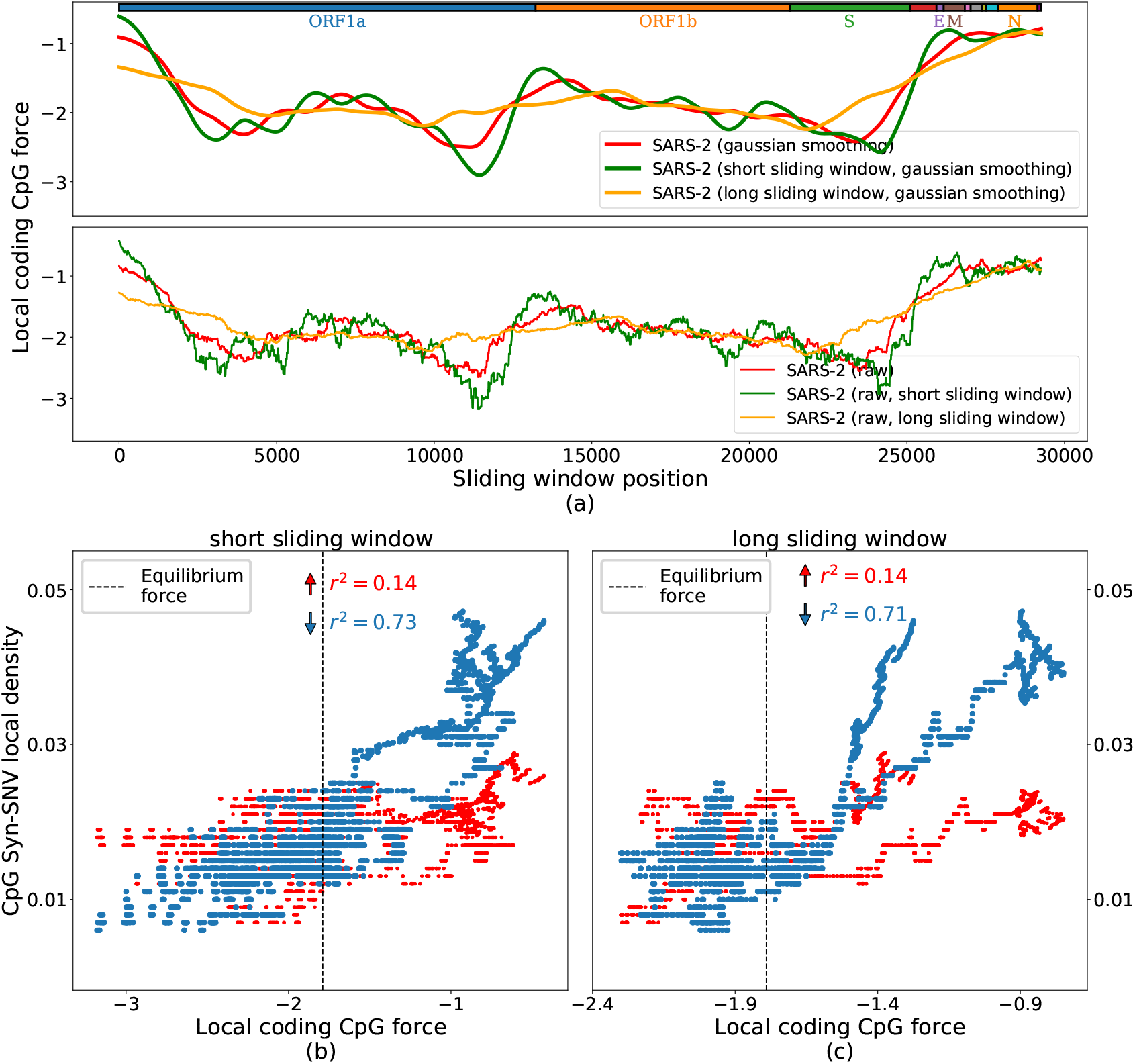
(a) Forces computed on SARS-CoV-2 ancestral genome, with different choices of sliding windows (upper subpanel), or without Gaussian smoothing (lower subpanel). The sliding windows used in panel (a) are: 3000 nt, 1500 nt (short sliding window), or 6000 nt (long sliding window). In panels (b), (c) we replicate the analysis performed for Fig. 4b with the short and long windows, to show that the results are qualitatively similar. The slightly lowest performance of the short sliding window is due to the high sensitivity of the short windows for local details of the sequence, while the large window analysis is plagued by finite size effects (right side of panel (c)), due to the fact that the size is too close to the total size of the sequence, together with the presence of many CpG motifs at the beginning and end of the sequence.

**Figure SI.10:**
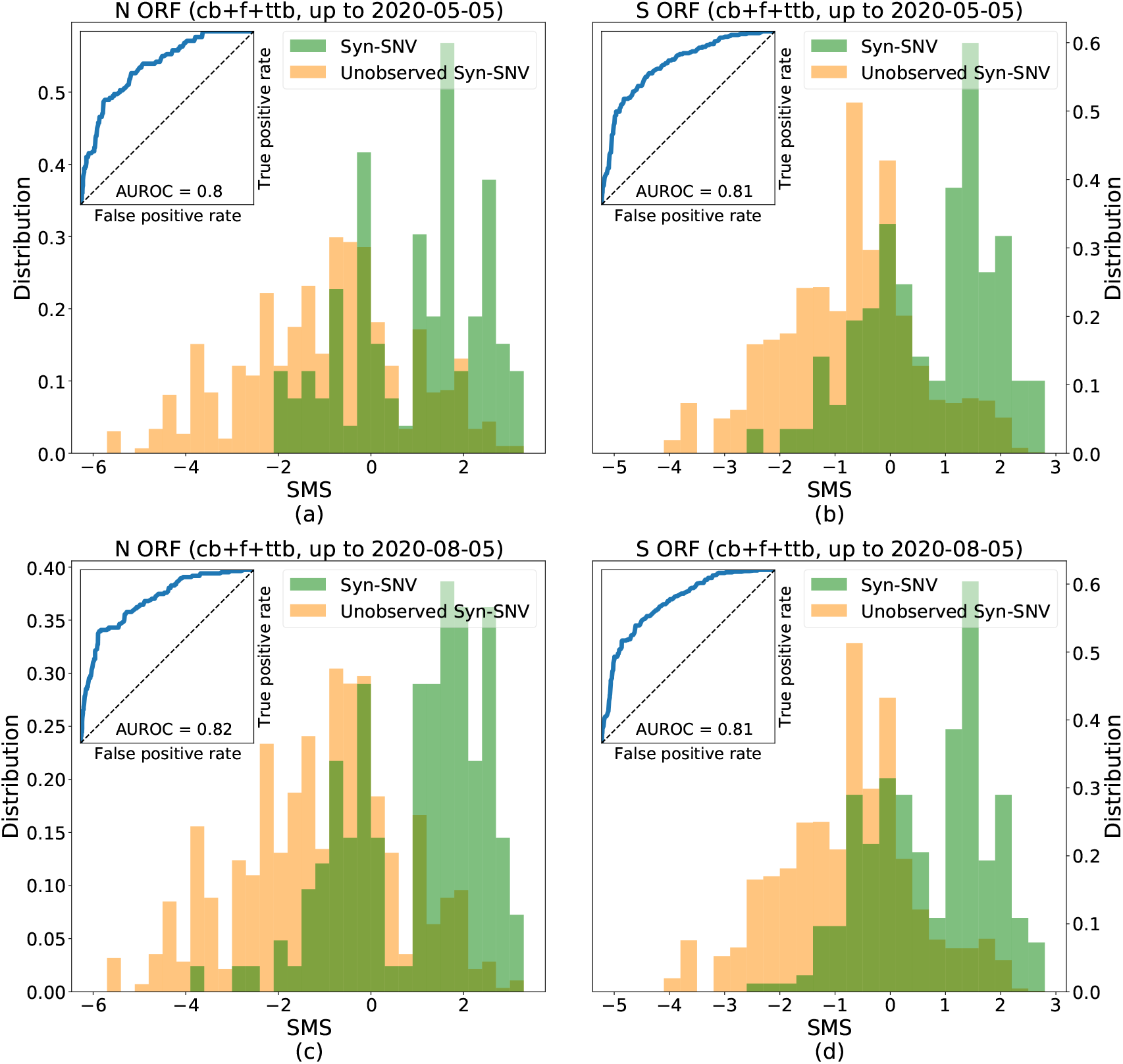
Same analysis performed in Figs. 5c and 5d, for sequences submitted to GISAID up to 5 May 2020 (panels (a), (b)) and up to 5 August 2020 (panels (c), (d)). The model cb+f+ttb can distinguish with large AUROCs (≥ 0.8), since 05 May 2020, observed Syn-SNV from conserved Syn-nt, and the model performance (intended here as AUROC) improves with time. Cutoffs for considering Syn-SNV are at 1 count for panels (a), (b), and at 2 counts for panels (c) and (d) (that is, approximately at 0.01% of the total number of collected sequences, as in main text for the full dataset). Other detailed analysis on sequences collected at previous times can be found in the several past versions of this manuscript on bioRxiv (https://www.biorxiv.org/content/10.1101/2020.05.06.074039v3.article-info).

**Figure SI.11:**
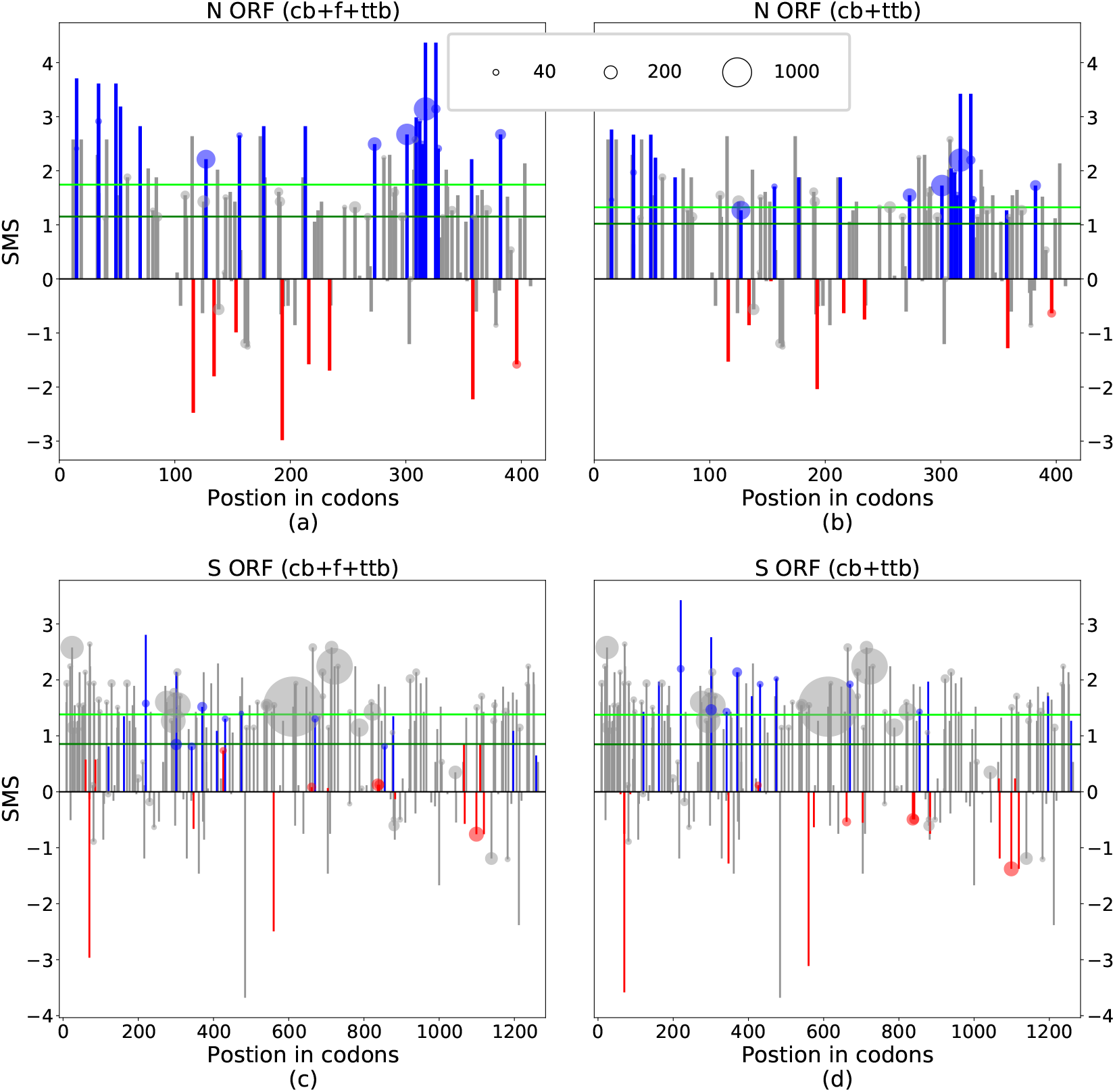
The SMS of all observed syn-SNV is given for N and S ORFs, with and without CpG drive. Bars colored in blue (red) correspond to CpG decreasing (increasing) syn-SNV. Circles on top of bars are drawn when the number of counts of the corresponding SNV is larger than 20, the size of the circle being proportional to the number of counts. Green (dark, light) horizontal lines give the average SMS (without or with counts).

**Table SI.1:**
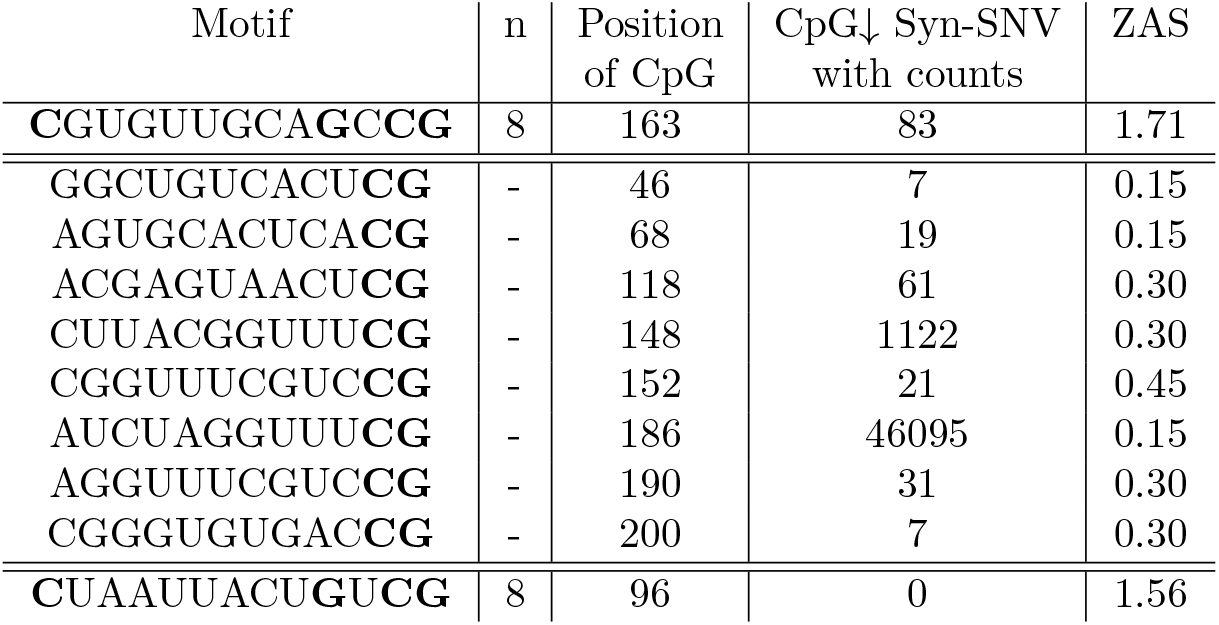
Analysis of CpG extended motifs of the form CnxGxCG with n = 4, 5, 6, 7 or 8 nucleotides in the 5’-UTR. The position is given with respect to the start of the 5’-UTR in the ancestral sequence, see Methods 4.6. All SNV happening in UTRs are considered synonymous. SNV observed with less than 5 counts are excluded from this analysis.

**Table SI.2:**
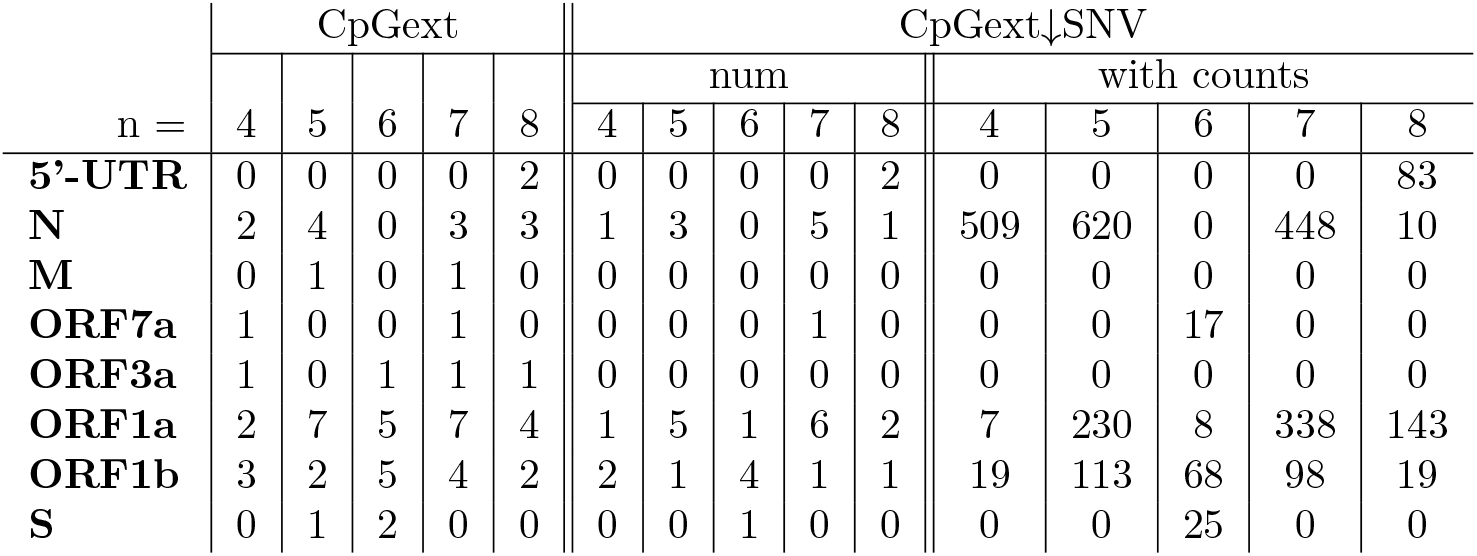
Supplementary information for Table 5. Only motifs of the form CnxGxCG with n = 4, 5, 6, 7 or 8 (CpGext) are considered and they are only characterized by the spacer lenght n. The number of CpGext motifs and of syn-SNV removing them is given for each value of n for each ORF or UTR (with at least 1 CpGext).

| n = | CpGext |  |  |  |  | CpGext↓SNV |  |  |  |  |  |  |  |  |  |
| --- | --- | --- | --- | --- | --- | --- | --- | --- | --- | --- | --- | --- | --- | --- | --- |
|  | 4 | 5 | 6 | 7 | 8 | num |  |  |  |  | with counts |  |  |  |  |
|  |  |  |  |  |  | 4 | 5 | 6 | 7 | 8 | 4 | 5 | 6 | 7 | 8 |
| **5'-UTR** | 0 | 0 | 0 | 0 | 2 | 0 | 0 | 0 | 0 | 2 | 0 | 0 | 0 | 0 | 83 |
| **N** | 2 | 4 | 0 | 3 | 3 | 1 | 3 | 0 | 5 | 1 | 509 | 620 | 0 | 448 | 10 |
| **M** | 0 | 1 | 0 | 1 | 0 | 0 | 0 | 0 | 0 | 0 | 0 | 0 | 0 | 0 | 0 |
| **ORF7a** | 1 | 0 | 0 | 1 | 0 | 0 | 0 | 0 | 1 | 0 | 0 | 0 | 17 | 0 | 0 |
| **ORF3a** | 1 | 0 | 1 | 1 | 1 | 0 | 0 | 0 | 0 | 0 | 0 | 0 | 0 | 0 | 0 |
| **ORF1a** | 2 | 7 | 5 | 7 | 4 | 1 | 5 | 1 | 6 | 2 | 7 | 230 | 8 | 338 | 143 |
| **ORF1b** | 3 | 2 | 5 | 4 | 2 | 2 | 1 | 4 | 1 | 1 | 19 | 113 | 68 | 98 | 19 |
| **S** | 0 | 1 | 2 | 0 | 0 | 0 | 0 | 1 | 0 | 0 | 0 | 0 | 25 | 0 | 0 |

**Table SI.3:** Supplementary information for Table 6, where all the quantities are computed through the model cb+ttb (without CpG drive).

|  | Average SMS |  |  | AUROC | ANOVA F-test |  |
| --- | --- | --- | --- | --- | --- | --- |
|  | SNV | u-SNV | diff |  | F | num items |
| **3'-UTR** | 0.33 | -0.39 | 0.73 | 0.77 | 72.2 | 81 + 405 |
| **5'-UTR** | 0.57 | -0.34 | 0.91 | 0.83 | 81.9 | 56 + 577 |
| **N** | 1.02 | -0.81 | 1.83 | 0.83 | 162.0 | 114 + 977 |
| **M** | 1.12 | -0.70 | 1.81 | 0.85 | 94.5 | 58 + 557 |
| **ORF10** | 1.37 | -0.51 | 1.89 | 0.84 | 9.8 | 6 + 87 |
| **ORF7a** | 0.30 | -0.77 | 1.08 | 0.71 | 12.4 | 23 + 289 |
| **ORF8** | 0.77 | -0.84 | 1.62 | 0.83 | 36.8 | 25 + 266 |
| **ORF3a** | 1.06 | -0.80 | 1.86 | 0.85 | 96.9 | 54 + 642 |
| **ORF1a** | 0.93 | -0.89 | 1.83 | 0.86 | 1580.3 | 848 + 9966 |
| **ORF1b** | 0.78 | -0.87 | 1.63 | 0.82 | 624.9 | 432 + 6026 |
| **E** | 0.35 | -0.56 | 0.91 | 0.68 | 3.6 | 10 + 217 |
| **S** | 0.85 | -0.90 | 1.75 | 0.84 | 375.2 | 223 + 2924 |
| **ORF6** | 0.35 | -0.56 | 0.91 | 0.70 | 5.3 | 10 + 127 |
| **ORF7b** | 0.70 | -0.41 | 1.11 | 0.77 | 6.1 | 9 + 98 |

**Table SI.4:** Supplementary information for Table 6, where ave SMS diff and AUROC are given for the models cb+ttb and cb+f+ttb while taking into account syn-SNV counts for the computation. In both cases, counts are used to weight observed syn-SNV. Notice that taking counts into account can lead to strong bias effects if few mutations are observed extremely more often than all others. For instance, in ORF1a the mutation observed more often has alone more than 50% of the total number of counts for the ORF, hence it strongly influences all indicators in this table. Similar effects are seen for M, ORF8 and 5’-UTR.

| model: | ave SMS diff |  | AUROC |  |
| --- | --- | --- | --- | --- |
|  | cb+ttb | cb+f+ttb | cb+ttb | cb+f+ttb |
| **3'-UTR** | 0.75 | 1.08 | 0.79 | 0.84 |
| **5'-UTR** | 1.54 | 3.18 | 0.97 | 0.99 |
| **N** | 2.13 | 2.73 | 0.88 | 0.89 |
| **M** | 0.95 | 1.08 | 0.68 | 0.68 |
| ORF10 | 1.44 | 1.67 | 0.79 | 0.76 |
| **ORF7a** | 1.75 | 2.10 | 0.84 | 0.85 |
| **ORF8** | 1.72 | 1.89 | 0.85 | 0.85 |
| **ORF3a** | 2.19 | 2.38 | 0.91 | 0.91 |
| **ORF1a** | 2.17 | 2.16 | 0.91 | 0.91 |
| **ORF1b** | 1.92 | 1.92 | 0.90 | 0.90 |
| **E** | 1.10 | 1.59 | 0.76 | 0.76 |
| **S** | 2.28 | 2.14 | 0.91 | 0.92 |
| ORF6 | 0.72 | 0.73 | 0.69 | 0.69 |
| ORF7b | 1.29 | 1.30 | 0.84 | 0.84 |

### SI.3 From CpG force to CpG relative abundance

We want to show in which limit the CpG force (without codon constraints) is equivalent to the relative dinucleotide abundance [1], Eq. 4. We start from the partition function:

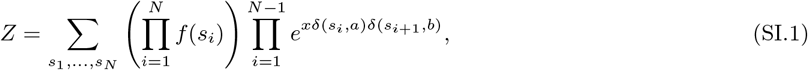

where *δ* denotes the Kroneker delta function. In the spirit of a cluster expansion, we write

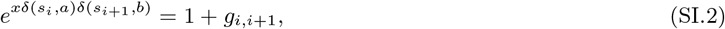

where

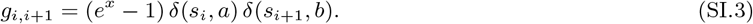

Inserting back this into Eq. (SI.1), we obtain

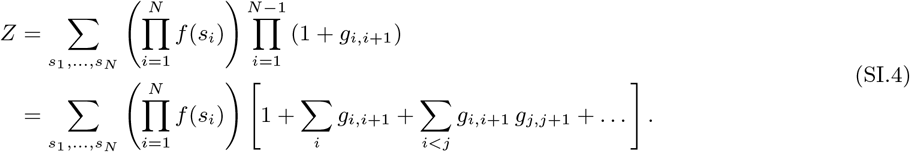

Now we can compute each term in the cluster expansion, and we get for the *k*-th term (for *a* ≠ *b*, as in the CpG case)

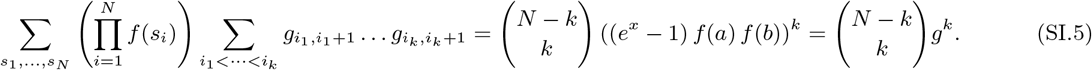

where we defined *g* = (*e^x^* − 1) *f*(*a*) *f*(*b*). Now we suppose *N* = 2*m*, that is *N* is even (however, we will consider soon the large-*N* limit, where this request is not necessary anymore). Therefore, we have

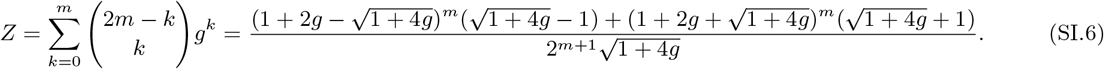

To proceed further, we can consider the case where *g «* 1. This is a good approximation when *x* ≃ 0, and it is also fairly good as long as *x* is lower than 0, but it is less good for the most negative forces observed here (see Fig. 3a).

Under this hypothesis, we have

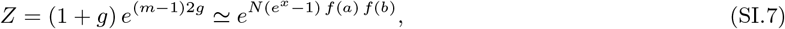

where in the last step we used also that *N* » 1. From this, by using that 〈*n*〉 = *∂x* log *Z* and requesting 〈*n*〉 = *n*_0_ = *Nf*(*ab*), we obtain Eq. 4. Fig. SI.12 shows the correlation between the CpG force with the nucleotide bias and the CpG relative abundance.

**Figure SI.12:**
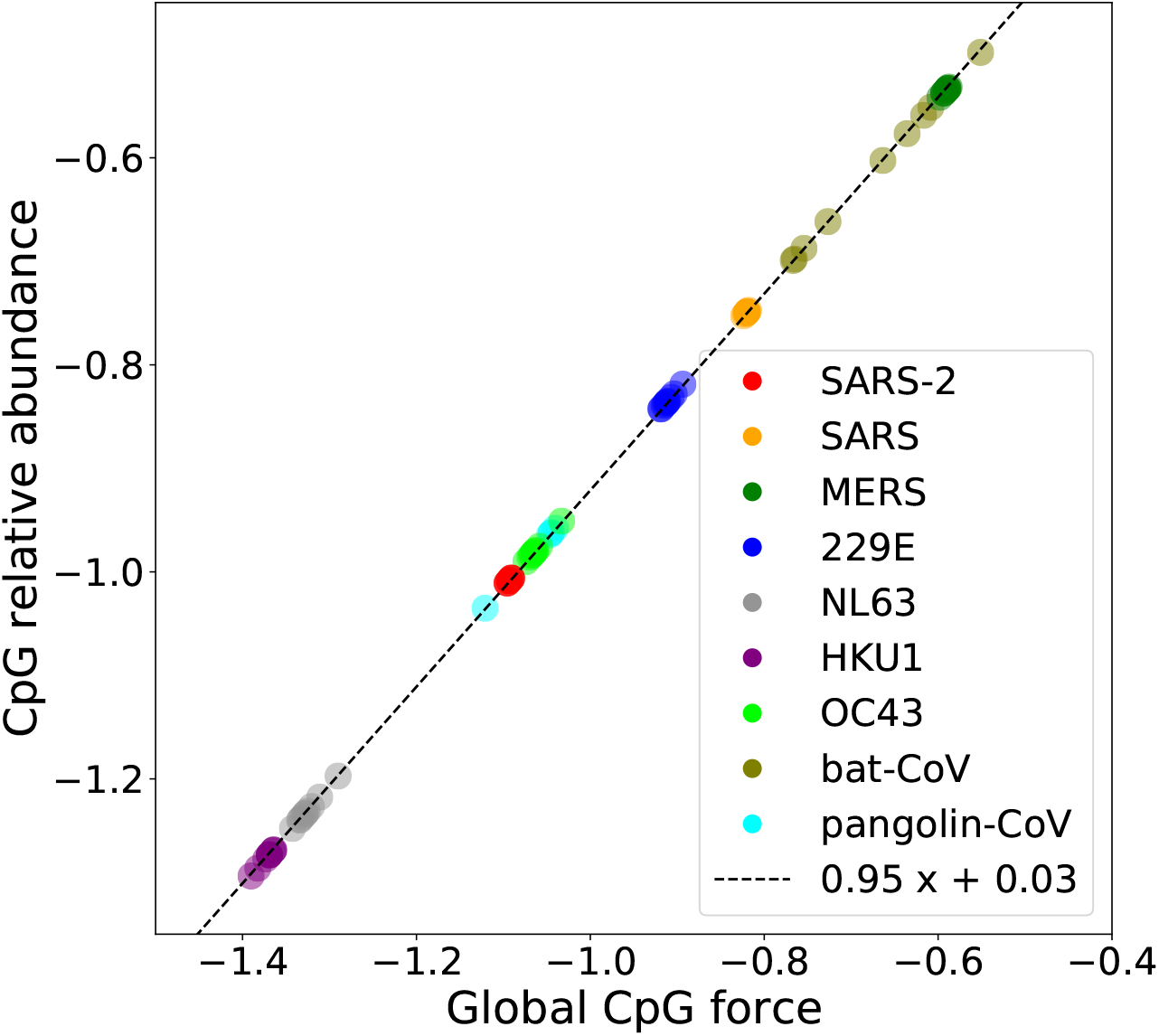
Comparison between the CpG force and the CpG relative abundance index. As discussed in Sec. SI.3, these two quantities are almost identical when the genome is long and the force is not too large in absolute value. Here 10 different genomes for several coronavirus species are used to compute these two quantities, and the dashed black line is a linear fit of the resulting points.

## Footnotes

1 Comparisons based on dinucleotide number rather than force give qualitatively similar results, see Suppl. Fig. SI.2.

2 Such cutoff removes very rare mutations which may be due to sequencing errors effect.

3 We consider here the virus codon bias, calculated on the Wuhan ancestral strain, rather than the human codon bias, as SARS-CoV-2 is likely not in equilibrium with its host yet. This choice will be justified later.

4 We consider the canonical ratio 4:1 here, see Methods Sec. 4.3.1 for details.

5 Here we drop the subscripts *nc* and *c* used in the previous section to identify non-coding and coding forces, since the SMS is defined for a generic force.

